# Ultra-slow conformational dynamics and catch bond formation of a Bacterial Adhesin revealed by a single-domain variant of FimH

**DOI:** 10.1101/2025.07.17.665197

**Authors:** Pearl Magala, Lisa M. Tuttle, Gianluca Interlandi, Laura A. Carlucci, Molly Y. Mollica, Maria K. Janowska, Wendy E. Thomas, Evgeni V. Sokurenko, Rachel E. Klevit

## Abstract

Bacterial adhesins such as FimH are critical for host colonization and persistence under the mechanical forces encountered at sites of infection such as the urinary tract. Despite decades of research, the molecular mechanisms by which FimH—a key virulence factor of *Escherichia coli*—regulates its binding through conformational switching remain incompletely understood. FimH operates across a range of conformations that includes low- (LAS), intermediate-, high-affinity (HAS) states-- and forms catch bonds which paradoxically strengthen under force. The allosteric pathways governing these transitions remain poorly defined due to experimental limitations that restrict understanding of key dynamic phenomena that underlie ligand-triggered conformational shifts and force-induced long-lived interactions. Such understanding is central to drug discovery efforts to target bacterial adhesion. Here we present a model system that fully recapitulates the conformational repertoire of FimH in the absence of its pilin domain. Our findings demonstrate that a single mutation in the lectin domain induces the LAS while allowing for ligand-binding induced conformational change to the HAS and catch bond formation, mirroring the behavior of the native FimH adhesin. We propose a dynamic allosteric mechanism that involves ultra-slow, low-frequency dynamics for the ability of FimH and the bacteria that express it, to sustain long-lived interactions with mannose under both static and force conditions.

**Significance:** Urinary tract infections (UTIs) are among the most common bacterial infections, and their initiation depends on the ability of uropathogenic *Escherichia coli* (UPEC) to adhere to bladder cells. The adhesion is mediated by FimH, a protein on the outside of UPEC that binds mannose-containing glycoprotein receptors and strengthens its grip under shear stress via a catch-bond mechanism. To investigate FimH function, we engineered a variant that can adopt both low- and high-affinity states of FimH and can form catch bonds. We discovered that FimH is governed by ultra-slow conformational dynamics that vary even among structurally similar states. These findings provide a mechanistic framework for developing anti-adhesive therapies that target FimH dynamics, offering a novel strategy to prevent and treat UTIs.

## Introduction

The urinary tract, a hostile environment of constant fluid flow, poses a challenge for bacterial adhesion and colonization. Yet, *Escherichia coli*—the leading cause of urinary tract infections (UTIs) (1, 2)—thrives in this dynamic setting, due largely to the conformational capabilities of its molecular adhesin, FimH (3, 4). Located at the tip of type 1 fimbriae, FimH acts as a molecular switch, converting among low, intermediate, and high-affinity states (termed LAS, IAS, and HAS, respectively) that allow it to grip or release mannose moieties present on the surface of host cells. Its so-called catch bond behavior, in which binding is stronger and longer-lived under shear forces such as urine flow, is essential for bacterial survival and virulence (5–10). However, the molecular mechanisms underlying conformational dynamics and allosteric regulation of FimH remain poorly understood.

FimH has two domains: a lectin domain (LD) located at the tip and a pilin domain (PD) that connects FimH to a fimbria. The LD binds mannose via loops on the opposite end of the domain from the PD-contacting surface (Fig. 1) (11–14). The PD allosterically regulates mannose binding: when the lectin and pilin domains are in contact with each other, the LD adopts the LAS conformation and upon ligand binding, the loops close around mannose, resulting in the IAS. Separation of the two domains by mechanical or shear force triggers a conformational change in the LD associated with the HAS (Fig. 1A) (5). The ability of FimH to switch among these states enables colonization and survival of uropathogenic *E. coli* (UPEC) in the urinary tract by modulating adhesion and detachment under both static and flow conditions.

**Figure 1:**
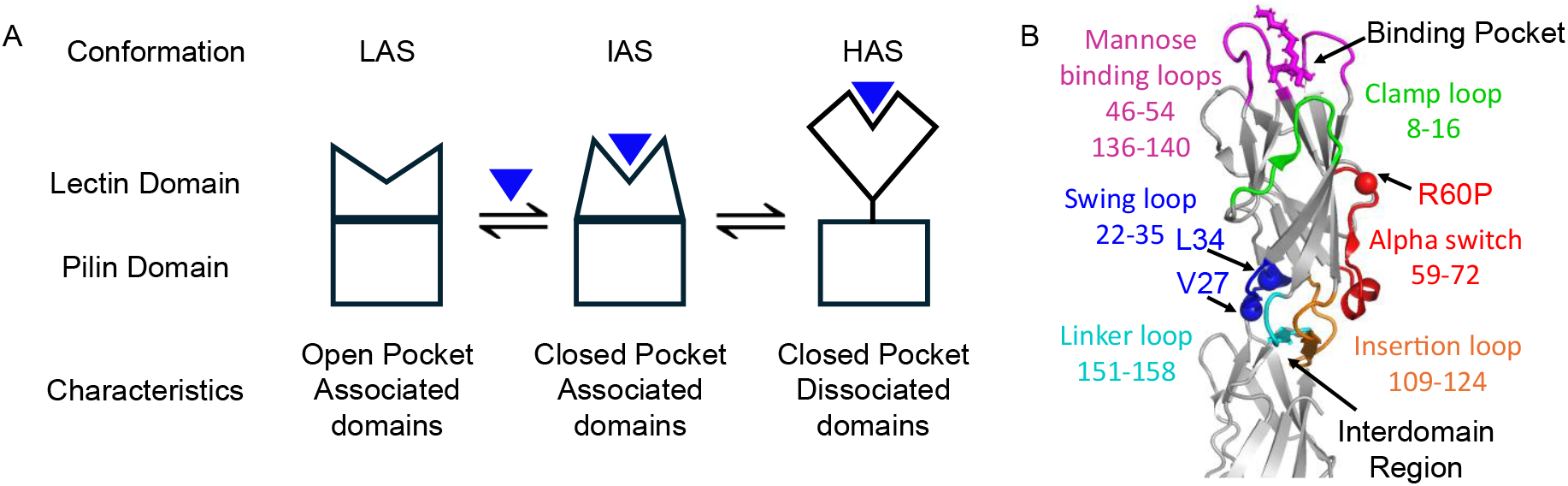
Conformational states in FimH. (A) FimH adopts LAS, IAS, and HAS conformations, characterized by: “open pocket and associated domains (LAS)”, “closed pocket and associated domains (IAS),” and “closed pocket and dissociated domains (HAS)”. (B) PyMOL representation of FimH^FL^ showing the binding pocket, interdomain region, regions that show the largest structural differences between HAS and LAS, and sites of mutagenesis (spheres) that allow LD to adopt LAS (PDB:4×0E).

Similar to force-induced domain separation, removal of the PD yields an isolated LD that spontaneously adopts the HAS conformation even in the absence of mannose. This behavior is consistent with a model in which contact between LD and PD stabilizes the LAS conformation (15, 16). Analysis of the many structures that have been solved of FimH reveals regions of the protein with the greatest structural differences between the LAS and HAS (Fig. 1B) (17–19). These regions have previously been targeted in attempts to produce LD variants that adopt the LAS in the absence of the pilin domain. A substitution in the Alpha switch region (residues 59-72) with R60P, or introduction of a disulfide bond within the Swing loop (residues 22-35), V27C-L34C (hereafter referred to as S-S) both adopt LAS in the absence of the PD, confirming that the LAS can be stabilized by forces other than contact with the PD (15, 20, 21). However, neither of these variants can convert to the HAS, signifying that they are conformationally and functionally “broken” even though neither mutant contains substitutions in either the mannose-binding loops or the PD-contacting loops.

We asked whether it is possible to design a version of the LD that both spontaneously adopts the LAS and retains its ability to transition to the HAS. Towards this end, we turned to residues in the LD that undergo large changes in their sidechain solvent accessibility between LAS and HAS conformations. Such residues were recently called “toggle switch” residues (22). We focussed on L34, whose sidechain is solvent-exposed in LAS and is core-buried in HAS (Fig. S1C) and discovered that substitution of Leu34 with lysine (L34K) creates a protein with the appropriate biochemical properties. The LD carrying L34K (“L34K^LD^”) adopts the LAS in the absence of the PD and mannose and transitions to the HAS upon mannose binding. Thus, L34K^LD^ represents a single-domain LD variant capable of recapitulating the behavior that underlies the function of FimH^FL^. The ability to create a single-domain allosteric model enabled the use of biochemical and biophysical approaches that are not possible with the full-length protein. We used a combination of biochemical binding experiments, Nuclear Magnetic Resonance (NMR) spectroscopy, Hydrogen-Deuterium Exchange (HDX), atomic force microscopy (AFM), and molecular dynamics (MD) simulations to investigate the functional, conformational, and dynamic properties of L34K^LD^. The results confirm that the single domain L34K^LD^ displays mannose-binding kinetics similar to the full-length protein and fully adopts the LAS and HAS conformations in the absence and presence of mannose, respectively. Finally, we demonstrate that bacteria expressing full-length FimH L34K retain their ability to aggregate red blood cells—a property that reflects catch bond formation. Altogether, the results elucidate an allosteric network that enables communication between the mannose-binding pocket at one end of the LD and the interdomain region at the other and, ultimately, the allosteric transition that gives rise to the enigmatic catch bond behavior.

## RESULTS

### Substitution of a toggle switch residue, L34, in FimH fimbriae inhibits transition to the HAS conformation

A monoclonal antibody that specifically recognizes FimH in the HAS conformation, mAb21, (23–26) was used to assess the ability of LD variants to adopt HAS in the context of fimbria. mAb21 binding to fimbria was measured in ELISA assays as a function of the FimH ligand, methyl α-D-mannopyranoside (hereafter referred to as mannose) concentration (Fig. 2 and Fig. S1B). WT FimH and a variant in which the L34 toggle residue was substituted with alanine showed similar mannose-dependent binding to mAb21, with little to no binding in the absence of mannose and increased binding with increased mannose concentration. This behavior is consistent with species that predominantly adopt a LAS conformation in the absence of mannose and adopt the HAS upon mannose binding. Variants that had L34 substituted with a charged sidechain (L34E and L34K) displayed low (L34E^LD^) or no (L34K^LD^) detectable mAb21 binding at the highest mannose concentration, indicating that these species do not adopt the HAS conformation recognized by the antibody under assay conditions.

**Figure 2:**
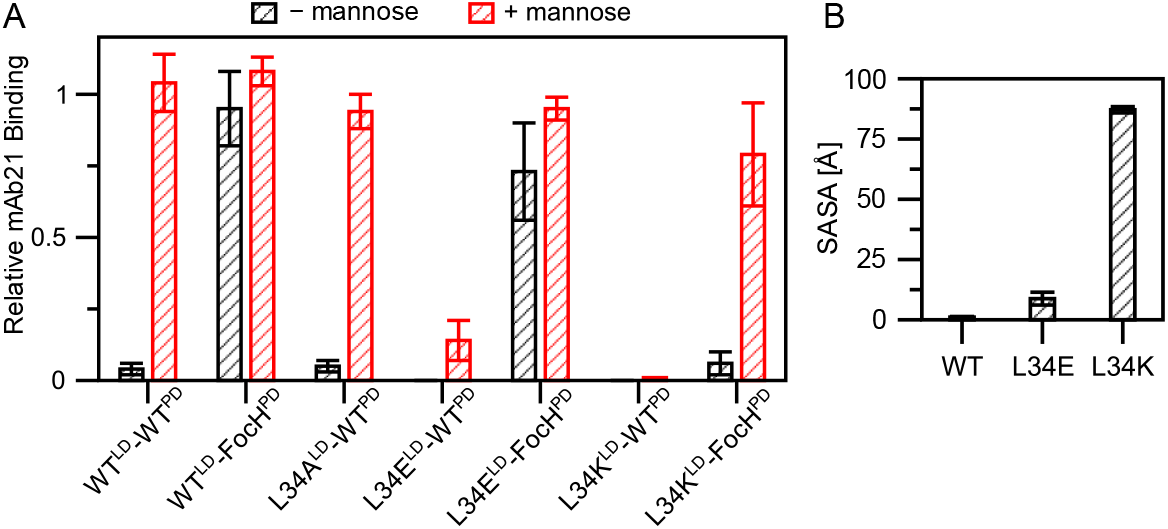
L34K disfavors HAS by favoring solvent exposure of the lysine sidechain. (A) Probing the HAS conformation with conformation-specific mAb21 binding. mAb21 binding was measured by ELISA on FimH variants incorporated into fimbria. Black bars, binding in the absence of mannose; red bars, binding in the presence of 2% mannose. Full binding curves are presented in Figure S1B. (B) Solvent accessibility of residue 34, averaged over the last 40 ns of a 50 ns MD simulation of the HAS LD.

The transition from LAS to HAS is allosteric and involves the mannose-binding pocket at one end of the LD and the interdomain region that makes contact with the PD at the other end (Fig. 1B). Mannose binding is activating (favors HAS) while PD binding is inhibitory (favors LAS). Thus, failure to adopt the HAS conformation upon mannose binding could be due to changes in either the mannose binding loops, the interdomain region, or both. To uncouple the two allosteric processes so that the intrinsic ability of LDs with toggle switch mutations to adopt HAS could be tested, we used a version of the PD called FocH (FocH^PD^) whose interdomain region is altered to make it incompatible with the LD and disrupting its ability to inhibit the HAS, (Fig. S1A) (6, 16, 27). In the FocH^PD^ background, WT^LD^ is fully bound by mAb21 in the absence of mannose, consistent with a loss of interdomain contact with the PD being sufficient to enable transition to HAS. L34E^LD^ also adopts HAS in the absence of mannose, as evidenced by the high level of mAb21 binding. In contrast, the L34K^LD^ chimera displays mannose-dependent binding behavior, consistent with its being in LAS in the absence of mannose and adopting HAS upon mannose binding (Fig. 2A and Fig. S1B). The results indicate that released or isolated L34K^LD^ retains the ability to populate HAS, but unlike the isolated/released WT^LD^, it requires mannose binding to do so. We note that the mAb21-binding experiments cannot distinguish whether L34K^LD^ adopts LAS or some other conformation in the absence of mannose and address this point below.

### Solvent accessibility of residue 34 in the isolated LD modulates LAS and HAS stability

The hydrophobic sidechain of toggle residue L34 is buried in HAS and is solvent-exposed in LAS. We hypothesized that substitution with a charged sidechain would favor the LAS conformation, where the sidechain faces solvent (Fig. S1C). The mAb21 binding experiments revealed that the lysine and glutamate substitutions are not equivalent, suggesting they are accommodated differently in the (buried) HAS conformation. To assess possible effects of substitution with Glu or Lys on the HAS conformation of the LD, molecular dynamics (MD) simulations of PD-free LD in the HAS conformation were performed and compared to simulations of WT^LD^ conducted in an earlier study (28). In the starting models, the E34 and K34 side chains faced into the protein core, as with L34. Solvent-accessible surface area (SASA) of the sidechains at position 34 averaged over the final 40 ns of 50-ns long simulations are compared in Figure 2B. The L34 side chain remained fully buried (SASA less than 1% of full exposure) (29). The glutamate side chain showed a modest increase in SASA, not significantly different from the wild-type L34 (∼5% of full solvent exposure; p-value > 0.1). However, the lysine residue averaged 41% of full exposure, significantly greater than that of both L34 and E34 (p-values < 0.05). Notably, the lysine side chain rotated from its core-facing orientation into the solvent within the first nanosecond of simulation time (Fig. S1D).

To assess how L34E and L34K might affect HAS stability, we conducted free energy perturbation calculations to estimate the change in free energy of folding due to a single-point mutation (ΔΔG). Calculations were performed for both the LAS and the HAS conformation of LD to evaluate destabilization in both states. The results indicate that L34K destabilizes HAS (where the sidechain is inward-facing) by a larger amount than it affects LAS. In contrast, L34E destabilized HAS by a similar amount as it destabilized LAS (Fig. S1E). Taken together, the simulations predict that glutamate at position 34 is relatively well tolerated in the buried orientation of the HAS conformation, whereas L34K^LD^ favors a conformation in which the lysine is solvent-facing—likely destabilizing HAS and consistent with the inability of L34K^LD^ to convert spontaneously to HAS in the absence of the PD.

### Similarity Analysis of NMR Spectra allows Conformational Classification of LD Variants

Structures of WT^LD^ with and without ligand and of R60P^LD^ in the absence of ligand have been previously determined (20, 30–32). WT^LD^ adopts HAS in both cases, and the structure of R60P^LD^ is identical to the LAS structure of the LD as observed in FimH^FL^ (C_α_ RMSD of less than 0.5 Å) (17). The (^1^H, ^15^N)-HSQC NMR spectrum of a protein provides a fingerprint that reflects its conformation, i.e., spectra of the same protein in two different conformations will differ. We therefore used NMR spectroscopy as a means to quickly ascertain conformations adopted by the position 34 variants using spectra of WT^LD^ and R60P^LD^ as reference states.

(^1^H, ^15^N)-HSQC NMR spectra were collected for each variant in the absence and presence of excess mannose. Visual inspection revealed striking similarities among certain spectra and large differences in others. For example, the spectra of WT^LD^ and L34A^LD^ in the absence of mannose are almost identical (Fig. S3) while the spectra of WT^LD^ and R60P^LD^ are markedly different (Fig. 3A). As well, spectra of R60P^LD^ and L34K^LD^ are highly similar (Fig. 3B). These qualitative comparisons suggest that L34A^LD^ is in the HAS (like WT^LD^) in the absence of ligand while L34K^LD^ is in LAS (like R60P^LD^).

**Figure 3:**
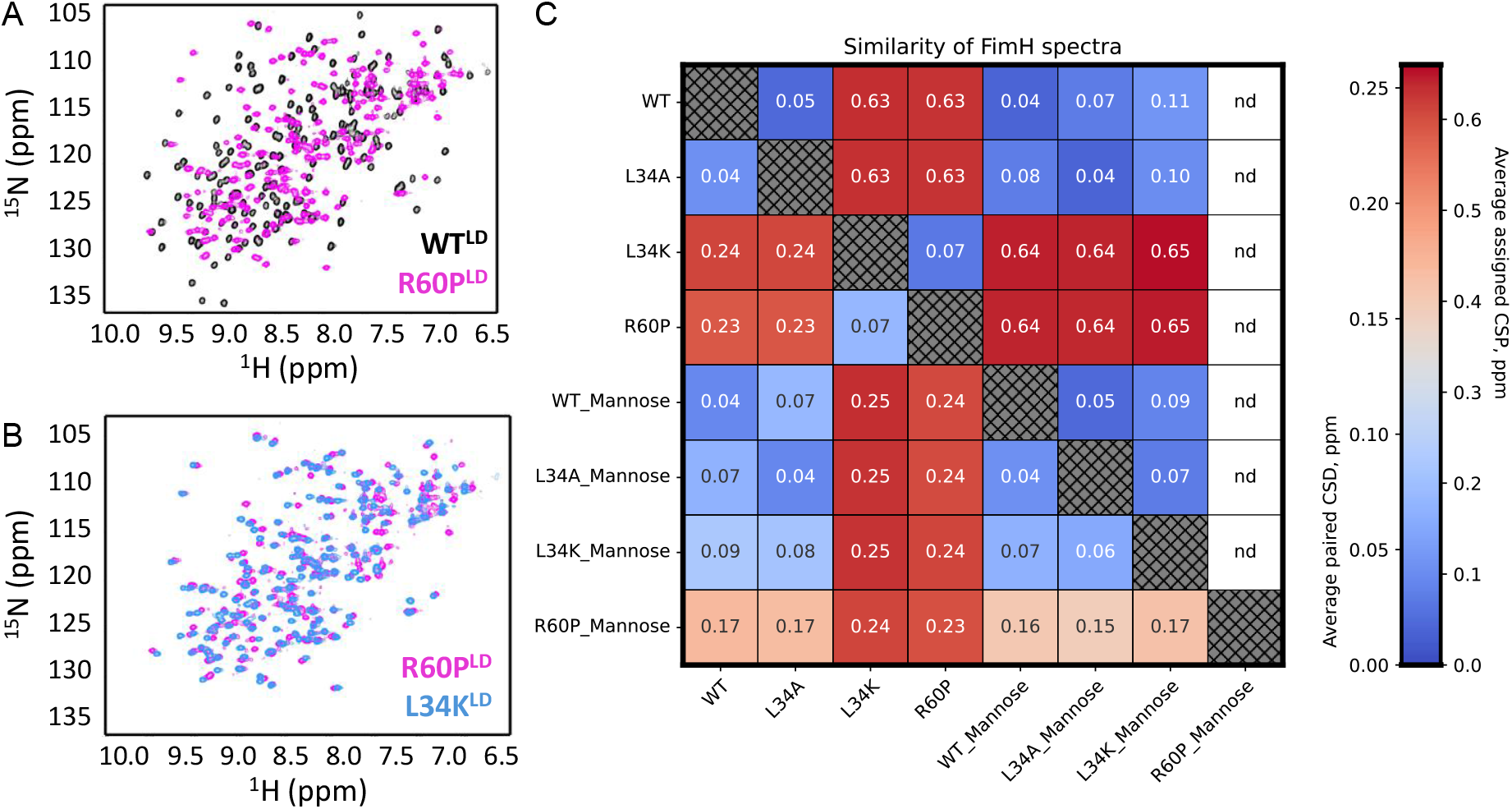
NMR Analysis of LD States reveals their conformations. (A) Overlay of the (^1^H,^15^N)-HSQC spectra of WT^LD^ and R60P^LD^ show that the spectra differ substantially, indicating distinct conformations. (B) Overlay of the (^1^H,^15^N)-HSQC spectra of R60P^LD^ and L34K^LD^ reveal that many residues overlap, suggesting that the conformations are the same. (C) Pairwise comparison of spectral similarity using the CSD score (bottom left) or assigned CSP (upper right; nd, not determined) reveals the similarity and differences between the LD variants in the presence and absence of mannose.

To build on these qualitative observations, we developed a quantitative analysis to assess the similarity of pairs of HSQC spectra. The peaks between two spectra are paired so that the summed total distance between peaks is minimized. A similarity score that corresponds to the average chemical shift distance (CSD) for all peaks is calculated. Because missing peaks can also be indicative of conformational differences between species, we assign a value of 0.5 ppm for each missing peak. Spectral pairs with the lowest score have the highest similarity (i.e., the peaks in that spectral pair overlap very well.) The CSD scores are presented in matrix format using heat map coloring in Figure 3C and Figure S2A (additional LD variants), with the highest similarity (lowest average chemical shift difference between two spectra) shown as blue and the lowest similarity shown as red. Consistent with conclusions made by visual inspection, the scores for WT^LD^ versus either R60P^LD^ or L34K^LD^ in the absence of mannose are the highest, indicating that the mutants adopt conformations different to that of WT^LD^.

Our analysis is agnostic to the true identity of each resonance in the spectra, raising a question as to its validity. The CSD score represents a lower bound on the chemical shift differences based on true peak identities. To validate the approach, we assigned all spectra except for R60P^LD^ in the presence of mannose, using standard (^13^C, ^15^N) 3D NMR experiments (33). The average chemical shift perturbation (CSP) of each pair of spectra was calculated from the assigned chemical shift positions and missing peaks were again assigned a value of 0.5 ppm. The CSP scores are presented in the upper right half of the matrix shown in Figure 3C and an example of the resultant peak pairings is shown in Figure S2C. As expected, the numerical CSP scores are higher than the CSD scores, but both the patterns and rankings remain the same, with the scores for WT^LD^ versus either R60P^LD^ or L34K^LD^ (in the absence of mannose) being the highest (∼0.65 ppm as opposed to ∼0.25 ppm using the assignment-agnostic approach). These results validate the use of the agnostic approach for comparing sets of HSQC spectra of protein variants, saving substantial time, effort, and resources required to assign NMR spectra. We refer to the analysis as “Chemical Shift Distance” (CSD) analysis and the resulting scores as CSD scores.

With few exceptions, spectra of LD variants fall into two categories: 1) high similarity to WT^LD^ and high dissimilarity to R60P^LD^ or 2) high dissimilarity to WT^LD^ and high similarity to R60P^LD^. Based on the known conformational states of WT^LD^ and R60P^LD^, the states of all other variants can be inferred, as shown in Table 1.

**TABLE 1.** Conformational States Adopted by Various LDs.

| Species | LD Conformational State Adopted |  |
| --- | --- | --- |
|  | No mannose | Mannose |
| FimH<sup>FL</sup> | Empty-LAS | Filled- IAS, HAS |
| R60P<sup>LD</sup> | Empty-LAS | Filled-Intermediate LAS |
| L34K<sup>LD</sup> | Empty-LAS | Filled-HAS |
| WT<sup>LD</sup> | Empty-HAS | Filled-HAS |
| L34A<sup>LD</sup> | Empty-HAS | Filled-HAS |

#### *WT*^*LD*^ *and* L34A^LD^ *adopt empty and filled HAS that differ only near the mannose-binding pocket*

As previously reported, WT^LD^ adopts a high-affinity state (HAS) and does not undergo extensive conformational changes when ligand is added (Fig. S4) (17, 18, 31, 34). The few chemical shift changes that occur upon ligand binding are indicative of local changes, with residues exhibiting significant chemical shift perturbations (CSP > 0.05 ppm), localized to the ligand-binding pocket, specifically, the two ligand-binding loops and the so-called clamp loop (Fig. S3). Thus, the NMR spectra of isolated WT^LD^ capture two conformations, namely, HAS with empty ligand-binding loops (“empty-HAS”) and HAS with mannose-bound ligand-binding loops (“filled-HAS”). Based on chemical shifts, the conformational difference between the two states is limited to the ligand-binding pocket. This behavior contrasts with that of the LD in FimH^FL^, which undergoes large conformational changes upon ligand binding, transitioning among LAS, IAS, and HAS.

#### R60P^LD^ adopts LAS but fails to achieve HAS

A crystal structure of isolated R60P^LD^ is identical to the WT^LD^ when in the presence of the PD, indicating that R60P^LD^ adopts the LAS conformation (20). R60P^LD^ exhibits many chemical shift changes upon addition of mannose, showing a ligand-induced conformational change, with a CSD score for R60P^LD^ in the absence and presence of mannose of 0.23 ppm. But comparison of R60P^LD^ with WT^LD^ in the presence of mannose gives a CSD score of 0.16 ppm, in the middle of the observed range of 0.04 - 0.25 ppm for all pairs. As the R60P^LD^:mannose spectrum lacks resonance assignments, residues with CSPs were identified based on the absence of an overlapping peak (Fig. S5). This approach revealed several things. First, the resonances that shift with mannose in WT^LD^ overlap with resonances found in the R60P^LD^:mannose spectrum, implying that the binding loops adopt a similar “filled” conformation in R60P^LD^. However, there are many resonances that show no peak overlap between the two spectra. When mapped on the structure of WT^LD^:mannose, these residues form two patches (Fig. S5C): one contains residues that surround R60 and a second forms a ring around the end of the domain that is distal to the ligand-binding site, i.e., the interdomain region of the LD. From this analysis, we conclude that while the R60P^LD^ binds mannose similarly to the WT^LD^, the mutation does not allow for the allosteric conformational change at the interdomain region to occur. Thus, NMR-based CSD analysis reveals that R60P^LD^ adopts the LAS conformation but does not attain the HAS conformation upon binding mannose.

#### L34K^LD^ adopts both the LAS and HAS conformations

In the absence of ligand, the spectrum of L34K^LD^ closely matches that of LAS-forming R60P^LD^, with a CSD score of 0.07 ppm. Upon mannose binding, the L34K^LD^ NMR spectrum undergoes extensive chemical shift changes indicating a ligand-induced conformational change. A comparison of the L34K^LD^ and WT^LD^ spectra in the presence of mannose showed few chemical shift differences, with a CSD of 0.07 ppm, confirming that L34K^LD^ adopts the HAS upon mannose binding. The observed CSPs are adjacent to the site of mutation, suggesting that the differences arise from the amino acid substitution, rather than a different conformation (Fig. S6). As mentioned above, substitution of L34 with the smaller Ala sidechain failed to produce an LD that adopts LAS in the absence of mannose, indicating that size, hydrophilicity, and/or charge of the toggle sidechain contribute to stabilize the LAS. Thus, the NMR spectra reveal that L34K^LD^ adopts the empty-LAS and filled-HAS conformations, similar to the LD in the context of FimH^FL^.

In sum, comparison of the NMR spectra of four variant LDs allowed for assignment of the conformational states adopted by each. The spectra reveal four different conformers: empty-LAS, empty-HAS, filled-HAS, and an intermediate-filled LAS. Table 1 summarizes the conformers adopted by each FimH variant in the absence and presence of ligand. Importantly, only L34K^LD^ mimics the states and transitions known to occur in FimH^FL^.

### Toggle switch variant L34K^LD^ displays rapid association and dissociation on mannose

NMR experiments showed that WT^LD^ and L34K^LD^ adopt the same conformational endpoint upon binding mannose. To determine whether this structural similarity translates to comparable binding kinetics, we used biolayer interferometry (BLI). Biotinylated mannose was immobilized on a streptavidin biosensor, and the kinetics of association and dissociation were measured for each LD variant. Response curves for WT^LD^ and L34K^LD^ are shown in Figure 4; additional LD variants are shown in Figure S7. The LD variants displayed distinct BLI response curves during both the association and dissociation steps. WT^LD^ showed a slow association phase and a slow dissociation phase. In contrast, L34K^LD^ demonstrated a rapid initial response followed by a more gradual phase during both association and dissociation (Fig. 4A). These distinct response profiles reflect underlying differences in the ligand binding properties among the LD variants.

**Figure 4:**
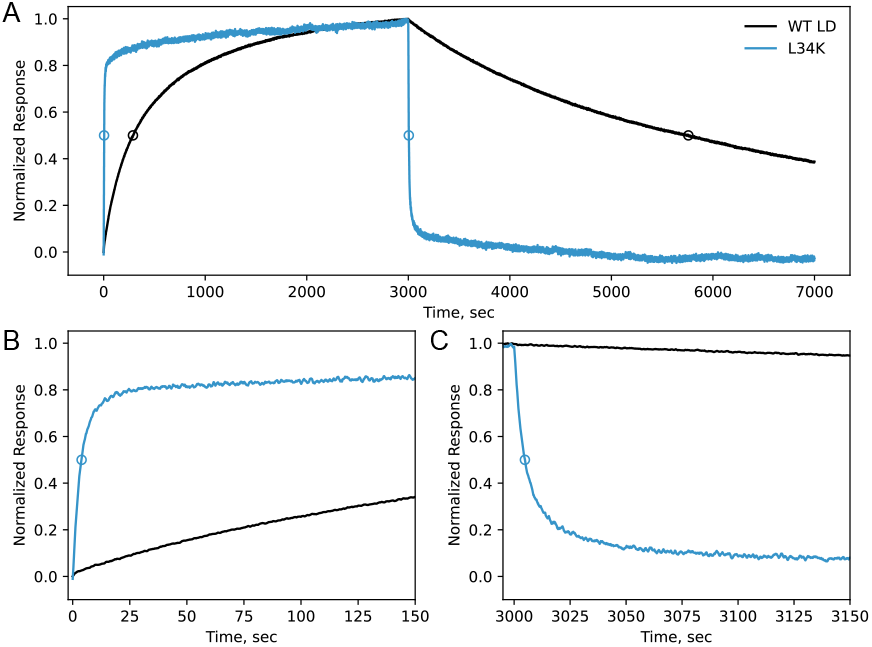
BLI response curves for WT^LD^ and L34K^LD^ reveal differences in binding kinetics. Response curves collected for 5 µM LD are shown. Black, WT^LD^; Blue, L34K^LD^. (A) The association phase covers the first 3000 sec, followed by the dissociation phase, from 3000 to 7000 sec. (B) and (C) show zoomed views of the first 150 sec of each phase. Circles indicate the t_1/2_ values as given in Supplemental Table S1 and Table 2.

BLI data were collected at five concentrations of each LD variant with serial dilutions starting from 5 µM; each gave high quality response curves. The data were fit to several models including a simple 1:1 binding model and a two-state binding model. No model was found to sufficiently capture all features of the curves. We therefore used a model-free empirical metric, the half-life (t_1/2_), where t_1/2_ is the time at which the measured response is 50% of the total response signal. The t_1/2_ for the association phase reflects both binding and dissociation events, while t_1/2_ for the dissociation phase reports solely on the dissociation process. Therefore, t_1/2, assoc_ values are concentration-dependent while t_1/2, dissoc_ values are not. Values are provided for the response curves collected at 5 µM LD and the average values for t_1/2, dissoc_ calculated from all curves are given in Supplemental Table S1.

The t_1/2, assoc_ for L34K^LD^, which is in the LAS at the beginning of the association phase is ∼70 times shorter than the WT^LD^ which is in the HAS conformation at the beginning of the association phase. The dissociation kinetics differ even more dramatically: the t_1/2, dissoc_ value for L34K^LD^ is more than ∼500-times shorter than WT^LD^. After five t_1/2_ time periods, a process will be ∼97% complete; we call this time period “T_assoc_” and “T_dissoc_”. The T_assoc_ values for L34K^LD^ and for WT^LD^ are 20 secs and 24 min, respectively. The difference in association time can be rationalized in part on the basis that the two variants start from different conformational states and in part because other processes including dissociation may occur during this phase. The dissociation phase reports solely on the dissociation of the LD from mannose and is more directly comparable. T_dissoc_ is 24 seconds for L34K^LD^ and almost 4 hours for WT^LD^. Importantly, dissociation occurs from the same conformational state, namely, filled-HAS (Table 1). This result strongly suggests there is something inherently different in the filled-HAS states.

The dramatic difference in kinetic behavior resembles the native-like behavior previously reported for two fimbrial forms of FimH (22). In one form, WT^LD^ is linked to the native WT^PD^ and is primarily in the LAS conformation in the absence of mannose. In the other, WT^LD^ is linked to the structurally altered FocH^PD^, with the LD predominantly stabilized in the HAS conformation. Although the association phases are not comparable due to differences in experimental setup, the dissociation phases can be meaningfully compared with our data (Table 2). The isolated LDs recapitulate the behavior of one of the full-length proteins: L34K^LD^ and the WT full-length protein (WT^LD^-WT^PD^) both dissociate rapidly, with T_dissoc_ values under 30 seconds, whereas WT^LD^ and the FocH chimera (WT^LD^-FocH^PD^) both dissociate very slowly, with T_dissoc_ values over 3 hours.

**Table 2.**
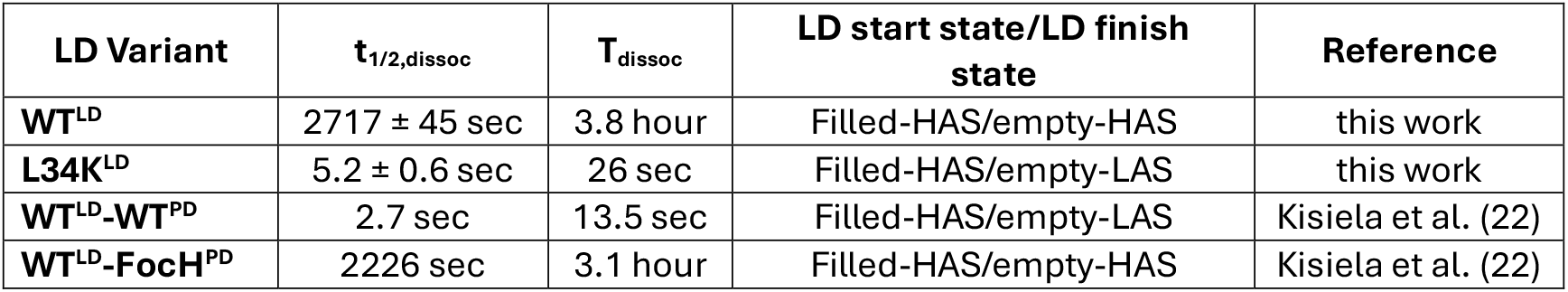
Dissociation times for Various FimH species.

The concurrence of the data is particularly noteworthy as the two experimental setups used different presentations of mannose (biotinylated mannose in the current work; RNaseB in Kisiela et al.(22)). In all cases, dissociation occurs from the filled-HAS LD conformation, but the end-point states differ, as indicated in Table 2 (LD start and finish states). Species that return to empty-LAS have short times to dissociate, regardless of whether the PD is present. Species that return to empty-HAS have very long dissociation times, again regardless of whether the PD is present. To summarize, the lectin domain with a single toggle residue substitution creates a single-domain module that recapitulates both the conformational and kinetic properties of the WT full-length adhesin protein.

### The FimH HAS exhibits very slow conformational dynamics

Although WT^LD^ and L34K^LD^ both adopt the HAS, there are very large differences in the rate at which they dissociate from mannose. Together, the observations suggest that the dynamic behaviors of the two variants may differ. Previous studies, including our own with WT^LD^, detected no remarkable dynamics in the nanosecond to picosecond timescale, as reported by NMR ^15^N T1 and T2 relaxation experiments (31). As allosteric motions are typically slower (milliseconds to seconds), we probed longer timescales ranging from milliseconds to hours. Attempts to observe dynamics on millisecond timescales reported by CPMG dispersion NMR experiments did not reveal any such dynamics. Extending towards even slower time regimes, we performed hydrogen-deuterium exchange mass spectrometry (HDX-MS) to investigate conformational dynamics (35). Ligand binding and conformational fluctuations can affect the solvent accessibility of backbone amides, influencing their deuterium uptake. Faster uptake indicates greater solvent exposure, while more buried amide groups and those involved in hydrogen bonds display slower deuterium uptake. Although we encountered some technical limitations (especially, incomplete coverage), the data revealed substantial differences among the HAS states of WT^LD^ and L34K^LD^ at long exchange times (30 minutes and longer), indicating that differences among the HAS species occur at very slow timescales (Fig. S8). Peptides exhibiting differences in deuterium uptake at long exchange times all included ligand-binding loop sequences (residues 3-21, 42-55, and 130-151).

To obtain more fine-grained information, we turned to HDX monitored by NMR, which provides residue-level information in the timeframe of minutes-to-hours. In our experiment, proteins lyophilized from H_2_O were dissolved in deuterium oxide buffer, quickly placed into an NMR spectrometer, and (^1^H,^15^N)-HSQC NMR spectra were collected continuously at five-minute intervals over a 20-hour period. H/D exchange was monitored from the intensity of each resonance as a function of time. Deuterium is silent in the NMR experiment, so exchange of an NH for ND causes loss of signal intensity that is directly related to the amount of deuterium uptake at a specific site.

Resonances exhibited three distinct exchange behaviors: 1) disappeared by the first spectrum (“rapid”, i.e., at 15 minutes they were fully deuterated; for example, blue traces for residue 27 Fig. 5A), 2) showed signal decay over the 20-hour time course (“intermediate”), and 3) maintained their intensity over 20 hours (“very slow”, for example blue and red traces for residue 11 Fig. 5A). A summary is provided in Supplementary Table S2. For residues with intermediate behavior, exchange rates (*k*_*ex*_) were determined by fitting decay curves to a single exponential model and half-life values were calculated (t_1/2, H/D_ = Ln(2)/*k*_ex_). For rapid residues, an upper value for t_1/2, H/D_ of 4 min was assigned; for very slow residues, a value for t_1/2, H/D_ of 270 hours was assigned. Residues with overlapping NMR signals were excluded from further analysis. Decay curves and exchange rate ratios and t_1/2, H/D_ are available as Supplemental Fig. S9.

**Figure 5:**
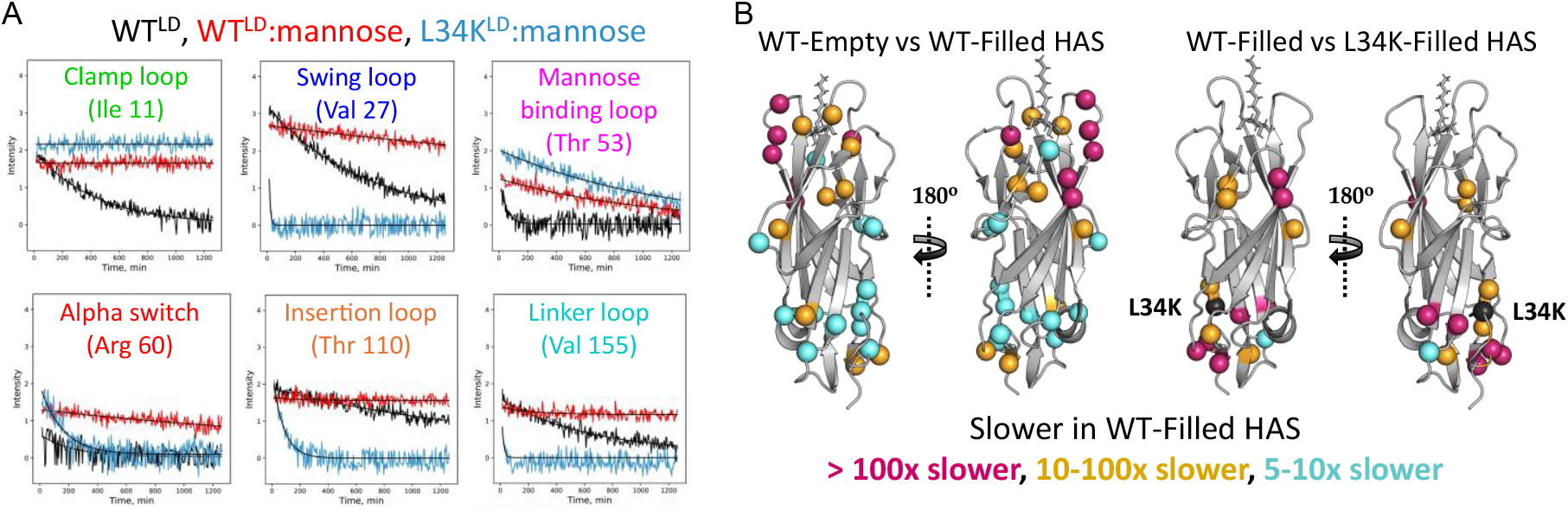
HDX/NMR reveals different dynamics for WT^LD^ and L34K^LD^ HAS states. (A) Exemplary HDX traces for resonances that correspond to the indicated regions. Black traces are for the empty-HAS; red traces are for WT filled HAS; blue traces are for L34K filled-HAS. Y-axis is peak intensity; x-axis is time. Traces that are flat and have intensity near/at zero are fully exchanged at the first time point; traces that are flat with intensity > 0 have very slow deuterium exchange. (B) Residues whose exchange rates differ by 5-fold or more between empty-HAS and filled-HAS are shown as spheres, with the image on the right rotated 180 degrees from the image on the left. Pink, residues with rates that are >100-fold slower in filled-HAS; Orange, residues with rates that are between 10- and 100-fold slower in filled-HAS; Blue, residues with rates that are between 5- and 10-fold slower in filled-HAS. The residues are 3, 5,11,13,15, 16, 27, 28,34, 35, 39, 46, 53, 60, 68, 72, 87, 95, 110, 112, 124,135, 138, 140, 141, 152, and 155. (C) Same as Panel B but for residues whose exchange rates differ by 5-fold or more between WT-filled HAS and L34K-filled HAS. L34 sidechain is shown in black spheres. These residues include 3, 10, 15, 16, 27-29, 32, 34, 35, 60, 72, 110, 112, 119, 152, and 155.

First, we compared apo-WT^LD^ and mannose-bound WT^LD^ to identify differences in conformational dynamics between empty-HAS and filled-HAS. Of the 111 resonances that could be compared across spectra, 30 have t_1/2, H/D_ values that differ by at least 5-fold between apo-WT^LD^ (empty-HAS) and bound WT^LD^, with six residues showing rate differences of >10^2^-fold (pink spheres in Fig. 5B). Those that slowed the most upon mannose binding are all within the mannose-binding loops and adjacent clamp loop. However, as is clear from Figure 5B, residues in more distal regions also display slower H/D exchange rates in the filled-HAS, with a large cluster of residues at the “bottom” of the LD. Most of the affected amides are observed to be hydrogen-bonded in solved structures of the filled-HAS conformation and should be resistant to H/D exchange. More rapid deuterium uptake indicates that these backbone groups visit exchange-competent environments (i.e., non-H-bonded) more frequently from empty-HAS than they do from filled-HAS. We note that these motions are very infrequent, with t_1/2, H/D_ values ranging from ∼4 hours to over 40 hours. Thus, while these regions are in the same conformation in empty- and filled-HAS, as evidenced by NMR chemical shift overlap, they transit away from their predominant conformation more often from empty-HAS. Mannose binding effects that are detected in residues at the other end of the LD provide direct experimental evidence of allosteric coupling between the mannose-binding pocket and the distal interdomain loops (Fig. 5B and 5C). Notably and unexpectedly, the allostery is manifested as differences in slow dynamics within conformations that differ only by whether they are ligand-free or ligand-bound. In other words, mannose binding to empty-HAS is associated both with protection of the mannose-binding loops (as expected) and a rigidification of the protein backbone at the opposite end of the domain that includes the PD-interacting loops.

Though the NMR spectra of L34K^LD^ and WT^LD^ are nearly identical in the presence of mannose, they exhibit dramatically different HD exchange behavior. Thirty-two resonances show much faster exchange kinetics in the L34K mutant. Surprisingly, no residues in the mannose-binding loops display different exchange rates, indicating that mannose affords similar protection in both cases. In contrast, residues in the adjacent clamp loop exchange between 10- and 10^3^-times faster in L34K^LD^, indicating that motions of this loop are not dictated solely by mannose binding. The L34K mutation also results in faster deuterium uptake in the swing loop, alpha switch, insertion loop, and linker loop (Fig. 1A and Fig. 5C). These findings further support direct allosteric coupling between the interdomain region where the mutation is located and the mannose-binding pocket, with the clamp loop being sensitive to both regions.

In conclusion, although allostery is typically observed as a conformational change in distal regions due to a binding event, the HDX data reveal allosteric effects that are manifested as changes in dynamics within a single conformation. Notably, the differences in HDX between apo and mannose-bound WT^LD^, and between bound WT^LD^ and bound L34K^LD^, occur in the same regions, suggesting that both mannose binding and the L34K mutation share a common allosteric mechanism.

### Toggle-switch mutant Fimbria can form catch bonds and mediate bacterial adhesion

Full-length WT FimH forms catch bonds with mannose that are characterized by an increasing fraction of long-lived, mechanically strong interactions under force. The model for catch bond formation by FimH invokes force-induced separation of its two domains. In this model, WT^LD^ starts in the LAS conformation in the absence of force (and mannose) due to its contact with the WT^PD^ and transitions to a state capable of forming catch bonds in the presence of mannose and force that separates the LD from the PD (8, 18, 36). The results presented above show that substitution of toggle residue L34 with Lys is sufficient to stabilize the LD in the LAS conformation, overcoming the requirement of PD binding at the interdomain interface (Fig. 2 and Table 1). Can an LD that adopts the LAS in the absence of the PD form catch bonds or is the act of domain separation *per se* required? We compared three species that each adopt the LAS conformation in the absence of mannose by Atomic Force Microscopy (AFM). Specifically, fimbria containing either 1) WT^LD^-WT^PD^, 2) L34K^LD^-FocH^PD^, or 3) S-S^LD^-FocH^PD^ at their tip were immobilized and a mannosylated BSA (man-BSA) coated cantilever was used to pull on individual fimbria (Fig. 6A). Sample force-extension curves are shown in Figure S10 for fimbriae with WT^LD^-WT^PD^ at the tip as a positive control, or no fimbriae as a negative control. Analysis of these curves confirmed that single bond ruptures are detected as force is applied in our experimental setup. Furthermore, the force-extension curves observed with S-S^LD^-FocH^PD^ or with L34K^LD^-FocH^PD^ were qualitatively similar, validating that we can compare the three species under the same set of AFM conditions.

**Figure 6.**
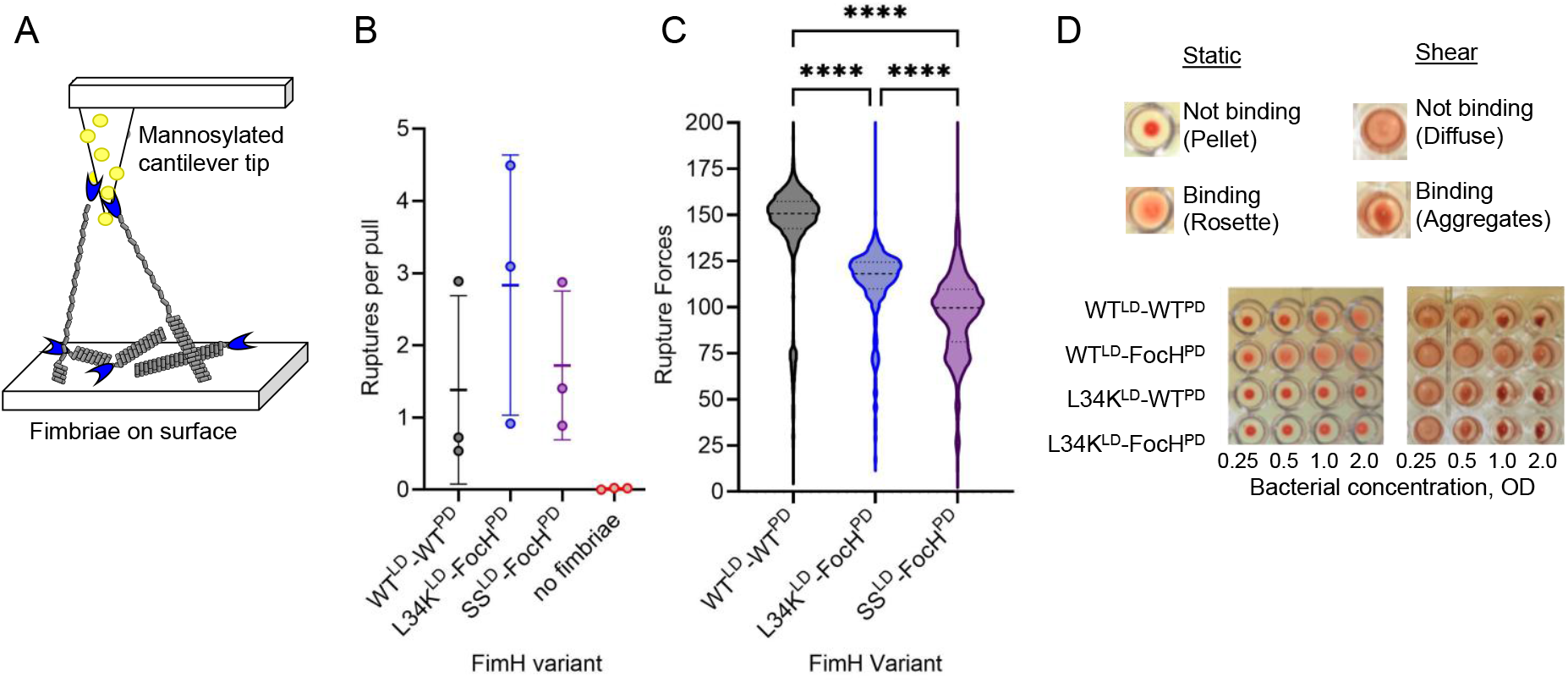
Measurement of the mechanical properties of FimH variants. (A) Schematic of the experimental configuration, with fimbriae immobilized on the surface, and mannose-BSA immobilized on the cantilever tip. (B) Mean number of ruptures per pull. Each dot indicates data from one of n = 3 experiments, and the error bars indicate standard deviation. (C) Violin plot of rupture forces for the three FimH variants. A one-way ANOVA was used to determine statistical significance with Tukey’s post-hoc analysis of multiple comparisons. *** indicates p < 0.0001. (D) Bacterial binding and catch bond formation assayed by the RBC assay under static and dynamic conditions.

Essentially no ruptures were observed in the no-fimbriae control (Fig. 6B, red) confirming that observed ruptures are specific to fimbriae. The number of ruptures per pull did not differ significantly among the three variants, indicating that the number of activated bonds generated under force is similar. As the man-BSA coated cantilever is pulled away from the surface to which the fimbriae are immobilized at constant velocity, a fimbria is stretched, leading first to its uncoiling (see Fig. 6A). This occurs at 75 ± 15 pN in our setup (Fig. S10).

Once a fimbria was fully uncoiled, the force was increased at a rate of several thousand pN/s until the FimH-mannose interaction ruptured, providing a measure of the rupture force of the bonds under a constant loading rate. Figure 6C shows a violin plot of the rupture forces observed for each species. WT^LD^-WT^PD^ maintains its association with the mannosylated tip at a high force (∼150 pN), with L34K^LD^-FocH^PD^ rupturing at slightly lower forces (∼120 pN). In contrast, S-S^LD^-FocH^PD^ ruptured at forces just slightly above those required to uncoil the fimbria to which it is attached. The results are consistent with the disulfide-bonded variant being unable to access the HAS conformation that enables long-lived associations under force. Furthermore, we can conclude that the toggle residue mutant can form long-lived associations under force, but its catch bond has slightly weaker mechanical strength.

Having observed the ability of fimbria containing the L34K toggle mutant to form long-lived interactions with mannose under mechanical force, we asked whether this property is retained in the native environment of bacteria. Bacterial strains expressing FimH composed of WT^LD^ or L34K^LD^ fused to either WT^PD^ or FocH^PD^ were tested for their ability to bind mannose moieties. Strains were tested for binding to N-linked high-mannose-type oligosaccharides of RNAse B on a surface (static conditions; Fig. S11). Binding was detected for each strain, confirming that functional FimH species are presented by each of the strains.

The ability of bacteria to bind to red blood cells (RBCs) under shear conditions results in the aggregation of RBCs and reflects their catch bond behavior (37, 38). Each strain was tested for its RBC binding under static and shear conditions. Under static conditions, binding to RBCs is manifested by bacterial-induced agglutination, resulting in the appearance of RBC “rosettes” at the bottom of U-shaped wells. (These appear as a diffuse pink area in the well). Over the course of an hour, rosette formation was observed for both WT^LD^-containing strains but for neither L34K-containing strain (Fig. 6D). This indicates that a WT-LD can transition to a conformational state capable of binding and remaining bound to the surface of RBCs under static conditions, regardless of whether its starting conformation is LAS or HAS (i.e., in the context of WT^PD^ or FocH^PD^, respectively). In contrast, stabilization of the LAS brought about by the toggle switch mutation creates a species that does not transition to a conformation capable of prolonged binding to RBCs under static conditions, consistent with its LAS conformation.

Strikingly different results were observed under shear conditions. Samples were subjected to continuous shaking for 20 minutes immediately after mixing bacteria and RBCs and assessed for visible RBC aggregation (Fig. 6D). (These appear as irregular, dense, dark red patches in the well.) As previously reported, FimH is highly effective in this assay (37). In contrast to the static case, both of the L34K-containing species induced RBC aggregation as robustly and rapidly as WT^LD^-WT^PD^. Once formed, these aggregates remained stable even after shaking was stopped. Thus, consistent with the results from AFM, the toggle switch mutant forms long-lived interactions under force. Altogether, we conclude that the L34K mutant, which is stabilized in the LAS but can fully transition to the HAS, displays full catch-bond behavior. This, in turn, suggests that domain separation itself is not required for catch bond formation. Instead, tensile force transmitted across the FimH via the ligand-binding pocket may be sufficient to directly induce the LAS-to-HAS conformational transition in LD.

## Discussion

The distinguishing feature of the bacterial virulence factor FimH underlying its ability to infect the urinary tract is its formation of catch bonds. Catch bonds are non-covalent interactions that become longer-lived under mechanical or shear force (5, 8, 39, 40). In the case of FimH, catch bonds enable bacteria to remain attached to mannose moieties on the host cell surface in the presence of urinary flow (∼20 mL/sec), promoting urinary tract infections. Structural studies have revealed that the FimH protein has a conformational repertoire that includes low-, intermediate-, and high-affinity states under static conditions. A wealth of experimentally-determined FimH structures has revealed conformational differences among the states, but there are no atomic-level structures solved of FimH actively engaged in a catch bond (5, 17, 41). Furthermore, static snapshots cannot identify the mechanism that underlies catch bond formation (42). As the ability to inhibit FimH from forming catch bonds is an attractive target for novel therapeutic strategies, we aimed to uncover the allosteric mechanism that regulates FimH and its ability to form catch bonds.

The business end of FimH is its lectin domain (LD), which binds mannose via two mannose-binding loops. Mannose binding is inhibited allosterically by the pilin domain (PD), which is also responsible for attaching FimH to fimbriae on the outside of bacteria. Contacts between the PD and LD maintain the LD in its low-affinity state, where the β-sandwich structure is short and wide and the binding pocket is shallow or ill-formed. Disruption of the interdomain interaction or removal of the PD favors adoption of the empty-HAS conformation, in which the β-sandwich structure is elongated (long and narrow) (16, 18). Binding of mannose leads to a closed pocket, but the filled-HAS conformation is otherwise not detectably different from the empty-HAS. The ability to transition between states is critical for bacterial adhesion to cell surface mannose moieties. The LAS allows for rapid association but also for rapid dissociation—a property that inhibits prolonged bacterial binding to free mannose moieties likely to be encountered physiologically (5, 43). The empty-HAS associates much more slowly due to its more closed binding pocket, but once bound, the filled-HAS dissociates from mannose on the order of minutes-to-hours. In the two-domain FimH protein, the PD plays an integral role in determining association/disassociation rates because it is required for adoption of the LAS (5, 20, 22, 43).

To understand the mechanics and properties that underlie the vastly different binding kinetics of FimH LD in its two end-states (i.e., LAS and HAS), we sought a mutation that would allow the isolated LD to spontaneously adopt LAS while retaining its ability to adopt HAS upon mannose binding. Previous attempts have successfully generated versions of the LD that spontaneously adopt LAS, but we show here that they do not transition fully to HAS. L34, a residue that was previously dubbed a “toggle switch” based on its sidechain orientation switching from outward-facing in LAS to inward-facing in HAS, was targeted with the notion that a charged residue at this position might favor LAS over HAS (22). Initial characterization of L34E and L34K mutants revealed that the two substitutions are not equivalent in their effects, as only L34K had the desired biochemical phenotype. The LD carrying L34K (“L34K^LD^”) adopts the LAS conformation in the absence of the PD and mannose but transitions to the HAS upon mannose binding. Thus, L34K^LD^ represents a single-domain LD variant capable of recapitulating the behavior that underlies the function of FimH^FL^.

Several previous studies have reported marked differences in mannose dissociation rates among variants of FimH. Although these studies differed in the specific form of mannose used, the experimental technique used, and the FimH species used, there is consensus that species that adopt the LAS in the absence of mannose dissociate much more rapidly (∼10^3^ – 10^5^ fold) than those that favor the HAS in the absence of mannose (5, 20, 22, 43). Measurements of binding kinetics of the isolated WT^LD^ and L34K^LD^ reported here reveal that they recapitulate both the association and dissociation behaviors of full-length proteins (Table 2). The result implies that the binding kinetics critical to FimH function is intrinsic to the LD. While a substantial difference in association kinetics can be rationalized on the basis that the two LD variants are in different starting conformations (LAS versus HAS), a similar explanation is not available for the >500-fold difference in measured dissociation, as both LDs assume HAS conformations that are virtually indistinguishable by NMR.

If the answer does not lie in structure *per se*, it must lie in the dynamics of the structure. Comparison of the H/D exchange rates of the two filled-HAS forms revealed that they differ in the rate at which backbone (amide) groups transit away from their predominant (mainly H-bonded) positions to ones in which they can exchange with solvent. If we consider these rare motions as a kind of “unsnapping,” then the backbone groups of filled WT^LD^ unsnap far less frequently than those of filled L34K^LD^. Notably, the sites where unsnapping occurs are the clamp loop that forms the side of the mannose binding pocket and the inter-domain region at the opposite end of the domain. The large effect of the toggle switch residue on the dynamics/rigidity of clamp loop residues that are 25 – 30 Å away suggests that this loop plays a major role in the allosteric mechanism. Although the clamp loop contains only one residue that contacts mannose, it assumes different positions in the LAS where it is swung open and the HAS where it is closed (18). The 10^2^-10^3^ -fold increase in H/D exchange of the clamp loop backbone in L34K^LD^ suggests a model in which rare motions of the clamp loop are coupled to the motions/rigidity of the inter-domain region. The burial of a lysine at position 34 in the HAS is met with increased dynamics of that region and these are transmitted to the clamp loop. Thus, in addition to the more canonical structural allostery inherent in the role of the PD in favoring the LAS over HAS, we propose that dynamic allostery (44, 45) within the LD in HAS itself is responsible for modulating the lifetime of a FimH-mannose interaction. This model is consistent with the dynamic allosteric model exemplified by Newton’s cradle, as suggested by Peacock and Komives (45). In sum, we propose that the lifetime of a HAS-mannose interaction under static conditions is dictated by rare “unsnapping” events in the clamp loop that allow for small movements of the mannose binding loops that enable mannose to leave the pocket. A corollary to this proposal is that FimH binders such as small molecules, antibodies, or other proteins that target the clamp loop and alter its dynamics will modulate the rate at which FimH dissociates from mannose moieties.

Despite the large difference in dissociation rate measured for WT^LD^ and L34K^LD^ under static conditions, both can form long-lived interactions under mechanical force (AFM) and catch bonds as reflected in RBC aggregation under shear conditions (Fig. 6). Without the ability to define the structure of an LD under force at atomic-level detail, we can only speculate as to the mechanism. It is possible that the catch-bond conformation is a new force-induced structure that differs from filled-HAS and that both WT^LD^ and L34K^LD^ can adopt such a structure. Alternatively, the catch-bond conformer may resemble filled-HAS but have even slower backbone dynamics that allow for prolonged interaction under force. While we cannot rule out the first option, we favor the second. We have identified the clamp loop’s dynamics as playing a key role in determining the rate of mannose dissociation. In the HAS, the clamp loop emanates from two anti-parallel β-strands (residues 7-12 and residues 15-21; Figure S12) both of which are H-bonded to a neighboring strand. On one side, the neighboring strand contains residues 1 – 5, where residue Phe1 makes direct contact with mannose. On the other side, the neighboring strand is the long, final strand of the LD (residues 143-157) that leads all the way from the mannose-binding pocket to the PD. A force that pulls the PD away from the LD will pull on this strand. One can imagine that a force that separates the PD from an LD that is bound to mannose on a cell surface will serve to lengthen the LD along that axis. Such lengthening would, in turn, result in neighboring atoms coming closer to each other. In model systems, the application of pressure (leading to compression and a shortening of inter-atomic distances) is associated with the formation of strong H-bonds from weaker ones (46). In the context of our unsnapping model, H-bonds would break even less frequently, leading to longer-lived complexes under force.

In summary, characterization of a previously proposed “toggle” residue that switches between outward-facing and inward-facing in the two end-point conformations of an allosterically regulated adhesin demonstrates the potential of designing protein constructs to mimic complicated behaviors (22). We found that a single switch mutation tips the balance of the LD to its low-affinity conformation, overriding the requirement of the allosteric effector, PD. The L34K^LD^ is able to form catch bonds—a distinguishing feature of FimH, revealing that catch-bond formation does not require active separation of the PD from the LD by force but, rather, is an intrinsic property of the LD itself under force. This approach has applicability beyond FimH and bacterial adhesins and could be extended to other adhesion proteins, including cell attachment proteins and viral attachment proteins, which play key roles in pathogenesis and cellular communication. Such information is crucial not only for understanding the mechanisms that underlie function but also for the design of effective inhibitors of the key conformational state(s) that allow bacteria to colonize and survive in a harsh host environment.

## Materials and Methods

### Generation of LD Variants by Site-Directed Mutagenesis

The *FimH* LD gene, cloned into the pET-22b plasmid, was subjected to site-directed mutagenesis using the QuickChange PCR-based method (47). Specifically designed primers were used to introduce the desired mutations. PCR amplification was performed with Phusion High-Fidelity DNA Polymerase under standard cycling parameters, followed by DpnI digestion to remove the original plasmid template. The resulting mutated plasmids were then transformed into *E. coli* Top10 cells, and colonies carrying the desired mutations were identified via sequencing. The verified plasmids were transformed into *E. coli* BL21(DE3) cells for protein expression.

### Protein Expression and Purification

#### Fimbriae

Bacteria expressing fimbriae were grown in 1 L cultures of LB broth supplemented with ampicillin and chloramphenicol, using 20 mL of an overnight starter culture for inoculation. The cultures were incubated at 37 °C while shaking at 125 rpm. Cells were harvested by centrifugation at 6000 rpm for 15 minutes at 4 °C and resuspended in 50 mM Tris-HCl, 150 mM NaCl, pH 7.4 (buffer A). Cell lysis was performed using an Osterizer blender, with five cycles of 1-minute blending followed by 1-minute rest intervals on ice. Cell debris was removed by centrifugation at 12,000 rpm for 15 minutes at 4 °C. Fimbriae were purified by precipitation with 2 M MgCl_2_ for 2 hours at room temperature under continuous stirring. The precipitate, containing fimbriae, was collected by centrifugation at 15,000 rpm for 30 minutes at 4 °C and resuspended in buffer A. Fimbriae concentration was determined using the bicinchoninic acid (BCA) assay (48).

#### LDs

The LD plasmid, containing a gene for ampicillin resistance and a His-tag, was transformed into *E. coli* BL21 cells and grown overnight at 37 °C on LB agar plates. The resulting colonies were used to inoculate 1 L LB cultures supplemented with ampicillin, and the cultures were grown at 37°C with shaking at 200 rpm until reaching an optical density at 600 nm (OD_600_) of 0.6–0.7. For ^15^N-labeling, the cells were grown in minimal media containing ^15^NH_4_Cl as the sole nitrogen source. The temperature was then reduced to 22°C for 1 hour prior to induction with 0.5 mM IPTG, followed by overnight incubation at 22°C. Cells were harvested by centrifugation at 6000 rpm for 15 minutes at 4 °C. Periplasmic extraction was performed using osmotic shock.

Specifically, the cell pellet was resuspended in 20 mM Tris-HCl, pH 8.0, containing 30% (w/v) sucrose (buffer B) and incubated at 4°C for 1 hour. Cells were pelleted by centrifugation at 12,000 rpm, resuspended in 20 mM Tris-HCl, pH 8.0 (buffer C), and incubated at 4 °C for 1 hour. The cell suspension was centrifuged at 16,000 rpm for 15 minutes, and the resulting supernatant was collected as the periplasmic extract. The lysate was applied to a gravity-flow nickel affinity column and washed with 20 mM NaH_2_PO_4_, 300 mM NaCl, pH 8.0 (buffer D). Bound protein was eluted with the same buffer containing 500 mM imidazole. The eluate was further purified by size-exclusion chromatography using a Superdex 75 column equilibrated with 20 mM Na_2_HPO_4_, 100 mM NaCl, and 0.5 µM EDTA pH6 (buffer E). The purified protein was concentrated using a 10 kDa molecular weight cutoff (MWCO) centrifugal filter at 4000 × g. Protein concentration was determined by measuring absorbance at 280 nm using a NanoDrop.

#### mAb21

mAb21 was generated from hybridoma cell cultures as previously described (26, 49) and purified using Protein G affinity chromatography. Briefly, supernatants containing secreted mAb21 were first clarified by centrifugation at 10,000 × g for 20 minutes and then filtered through a 0.22 µm membrane to remove cellular debris. The clarified supernatant was loaded onto a Protein G column pre-equilibrated with binding buffer consisting of 20 mM sodium phosphate, pH 7.0 (buffer F). After sample loading, the column was washed with 10 column volumes of the same buffer to remove unbound material. mAb21 was eluted with 100 mM glycine-HCl, pH 2.7, and eluted fractions were immediately neutralized with 1 M Tris-HCl, pH 9.0. The eluate was further purified by size-exclusion chromatography using a Superdex 200 column equilibrated with buffer F.

### ELISA experiments

To assess mAb21 binding, microtiter plate wells were coated with purified fimbriae at a concentration of 0.1 mg/mL in 0.02 M NaHCO_3_ buffer pH 7.0 (buffer G) and incubated for 1 hour at 37 °C in the absence or presence of mannose. After washing, mAb21 was added at a concentration of 0.5 μg/mL and incubated for 45 minutes. Bound antibodies were detected using a 1:5,000 dilution of HRP-conjugated goat anti-mouse Fc antibody. The reaction was developed at room temperature using 3,3′,5,5′-tetramethylbenzidine (TMB), and absorbance was measured at 650 nm.

### Molecular dynamics simulations

#### General setup and simulations with single-point mutations

The initial conformation was derived from the crystallographic structure of isolated LD in HAS with PDB code 1UWF (30). Single-point mutations, either L34E or L34K, were introduced by replacing the corresponding side chain in the wild-type and subsequently performing with the program CHARMM (50). 100 steps of steepest descent minimization in vacuo while the positions of all atoms except the mutated residue and residues within 4 Å were kept fixed. The simulation setup was similar as in a previous study (28). Briefly, the simulations were performed with the program NAMD (51) and the CHARMM22 force field (52). Each protein variant was solvated using a cubic water box with side length of 86 Å and periodic boundary conditions. Chloride and sodium ions were added to neutralize the system and approximate a salt concentration of 150 mM. During the simulations, the temperature was kept constant at 300 K by using the Langevin thermostat (53) with a damping coefficient of 1 ps^-1^, while the pressure was held constant at 1 atm by applying a pressure piston (54). Before production runs, harmonic restraints were applied to the positions of all heavy atoms of the protein to equilibrate the system at 300 K during a time length of 0.2 ns. Then, harmonic restraints were applied only to the heavy atoms of the mutated and the neighboring residues and the equilibration was continued for another 2 ns. After this equilibration phase, the harmonic restraints were released and two 50-ns long simulations were performed with each of the L34E and L34K variants. The simulations with the single-point mutations were compared with previously published runs performed with wild-type LD in HAS (28).

#### Free energy perturbation calculations to estimate the change in free energy of folding upon single-point mutations

To evaluate the change in free energy of folding due to single-point mutation in LD in both HAS and LAS, we made use of so called alchemical transformations (55), which were accomplished through free energy perturbation (FEP) calculations (56), similarly as in a previous study (57). Each alchemical transformation was performed in triplicate. The initial conformations for HAS were taken from snapshots along 50-ns long simulations performed in a previous study (57). To generate initial conformations for LAS, we performed two 50-ns long simulations starting with LD from the X-ray structure with PDB code 4XO9 (5) (only the first 160 N-terminal residues were kept). Each alchemical transformation was performed in the forward and backward direction. In the forward transformation, the wild-type side chain is slowly converted to the mutant side chain. The conformation achieved in the forward transformation is then used to start a backward transformation where the mutated side chain is converted back to the wild-type. During the process, the amount of work needed for each transformation is calculated. The forward and backward calculations were then combined and a value for the ΔG of the transformation was obtained using the Bennett’s acceptance ratio method (58) implemented in the ParseFEP plugin of VMD (59). Each forward and backward transformation was performed for 1 ns during which a parameter λ was varied from 0 (wild-type) to 1 (mutant) and from 1 to 0, respectively, in time intervals of the length of 0.025 ns for a total of 40 intermediate states. The first half of each time window involved equilibration and the second half data collection. Alchemical transformations were performed in the folded and in the unfolded state. The latter was approximated by the situation where residue 34 is fully solvent exposed, which was accomplished by using an Ala-Leu-Ala tri**-**peptide (a 50-ns long simulation was performed with Ala-Leu-Ala to sample initial conformations). The difference in the free energy of folding upon a single-point mutation can then be approximated by the difference between the ΔG values calculated from the alchemical simulations in the folded and unfolded state, respectively, similarly as in a previous study (57).

### NMR data acquisition, processing and analysis

(^1^H,^15^N)-HSQC experiments were acquired on a Bruker Avance III 600 MHz spectrometer equipped with a cryoprobe. Samples were prepared in 20 mM Na_2_HPO_4_, 100 mM NaCl, and 0.5 µM EDTA at pH 6 (buffer H). Backbone chemical shift assignments were obtained from standard triple-resonance experiments (HNCA and HNCACB) performed on a Bruker Avance III 800 MHz spectrometer equipped with a cryoprobe. All NMR experiments were conducted at 298 K. NMR data were processed using standard protocols in NMRPipe (60) and analyzed in NMRView (61).

For each sample, peaks in the (^1^H,^15^N)-HSQC spectra were picked using NMRViewJ at comparable levels. For spectra with known chemical shift assignments, the average amide chemical shift perturbation (CSP) is calculated according to Δδ_NH_ (ppm) = sqrt [Δδ_H_^2^ + (Δδ_N_/5)^2^]. An assignment free, pairwise spectral comparison is also possible. Considering each spectrum to contain peaks at coordinates (δ_H_, δ_N_/5), we calculate the matrix of all possible distances between peaks of one spectrum to the peaks in another spectrum. The pairwise similarity value – the chemical shift difference (CSD) score – is obtained from solving the two-dimensional rectangular assignment problem, wherein peaks are uniquely paired such that the sum of the distances between all peaks is minimized (62). We use the ‘linear_sum_assignment’ function in the SciPy optimize module to compute these values, which are normalized by the number of pairs (63). In cases where samples have different numbers of peaks, for both the CSP and CSD scores, missing peaks are considered to have a distance of 0.5 ppm.

#### HDX-NMR sample preparation and data collection

To prepare samples for HDX-NMR, purified protein was first dialyzed extensively against Milli-Q water and subsequently lyophilized. The resulting powder was reconstituted in a deuterium oxide (D_2_O)-based buffer containing 20 mM Na_2_HPO_4_, 100 mM NaCl, and 0.5 µM EDTA, adjusted to pD 5.6. NMR data acquisition was initiated 10 minutes after dissolution. A series of two-dimensional [^1^H,^15^N]-HSQC spectra were then recorded at 5-minute intervals over a 20-hour period to monitor D_2_O uptake.

### BLI data acquisition, processing and analysis

BLI streptavidin biosensors were hydrated in PBS buffer pH 7.2, containing 2% (w/v) BSA and 0.2% Tween-20 (buffer I). The proteins were also in the same buffer. Biotinylated mannose (0.05 ug/mL) was immobilized to the sensors for 600 s and then washed with buffer I for another 600 s. The sensors were then dipped in a serial dilution of proteins concentrations (5 µM – 0.312 µM) for 3000 s during the association phase. The dissociation phase was carried out by moving the sensors into buffer I for 4000 s. Data were collected on an Octet R8 instrument (Sartorius).

BLI data were exported from the Octet software for further analysis. The BLI data were collected at five concentrations with serial dilutions starting at 5 µM, each giving high quality response curves. These data were not sufficiently well-fit by several different models including a simple 1:1 binding model and a two-state binding model. We instead quantified ligand binding kinetics of the 5 µM traces using a model free empirical metric, the half-lives (t_1/2,on_ and t_1/2,off_), where t_1/2_ is the time at which the measured response is 50% of the total signal starting from either the beginning of the association or dissociation timepoints, respectively.

### HDX-MS data acquisition, processing and analysis

FimH LD proteins were exposed to an 85% D_2_O buffer (Buffer H) at room temperature for varying durations—4 seconds, 1 minute, 30 minutes, and 20 hours—to monitor hydrogen-deuterium exchange over time. To prepare fully deuterated controls, proteins were incubated in the same buffer at 75 °C for 30 minutes. The exchange reaction was halted by adding a chilled quenching solution (1.6% formic acid), followed by immediate flash freezing in liquid nitrogen. Frozen samples were later thawed and subjected to enzymatic digestion using immobilized NEP-2. The resulting peptides were analyzed with a Waters Synapt G2-Si mass spectrometer configured using a custom LEAP PAL automation system(64). Peptide identification was carried out through tandem MS on a Thermo Orbitrap Fusion Tribrid or via MSE on the Synapt G2-Si. Data were processed using either ProteinProspector (UCSF) or ProteinLynx Global SERVER (Waters). Deuterium incorporation levels were calculated with HDExaminer 3.0 (Sierra Analytics), and data analyses were performed using pyHXExpress(65).

### AFM data acquisition, processing and analysis

Type 1 pili expressing either WT^LD^-WT^PD^, L34K^LD^-FocH^PD^ and S-S^LD^-FocH^PD^ were immobilized at a concentration of 10–20 μg/mL in 0.02 M NaHCO_3_ buffer on 35 mm tissue culture dishes (Corning) for 75 minutes at 37 °C. Dishes were then washed three times with 0.2% bovine serum albumin in phosphate-buffered saline (PBS-BSA) and stored overnight at 4 °C in the same buffer to block nonspecific binding. For the negative control, the pili immobilization step was omitted. Olympus Biolever cantilevers were functionalized by incubation with 200 μg/mL mannosylated bovine serum albumin (man-BSA) for 75 minutes at 37 °C, followed by washing and overnight storage in 0.2% BSA-PBS at 4 °C. Force measurements were conducted using an Asylum MFP-3D Atomic Force Microscope. Cantilevers (spring constants of 4.38 and 6.36 pN/nm) were brought into contact with the sample surface to allow bonds to form between FimH and mannose, then retracted at a constant speed of 2 μm/s. Pulling experiments were performed in 0.2% BSA-PBS. Approximately 100 force curves (pulls) were collected at each of three to five different positions per condition on each day. Although multiple bonds often formed during each approach, the uncoiling of pili during retraction—combined with their variable lengths—resulted in well-separated rupture events, allowing clear resolution of individual bond rupture forces. Experiments were repeated on three separate days. Bond rupture forces were calculated using a custom automated script in Igor Pro 5.05A (WaveMetrics Inc., Lake Oswego, OR). Rupture force was defined as the difference between the force immediately before rupture and the average post-rupture baseline, where that difference is greater than four standard deviations in the force measurements in a region of constant force. Most nonspecific binding in the negative control occurred within 0.25 µM of the surface, so this was set as a minimum distance threshold to identify specific bonds.

### Bacterial adhesion assay

Bacteria were cultured overnight at 37 °C under static conditions. The following day, cultures were pelleted by centrifugation, washed twice with phosphate-buffered saline (PBS), and resuspended to an optical density of 2.0 at 540 nm (OD_540_). Flat-bottom 96-well plates were coated with 100 μl of bovine RNase B (20 μg/ml) in 0.02 M sodium bicarbonate buffer and incubated at 37 °C for 1 h. Wells were subsequently blocked with 0.2% bovine serum albumin (BSA) in PBS at 37 °C for 30 min and then aspirated. Control wells were coated with 0.2% BSA in PBS alone. Bacterial suspensions were added to the coated wells and incubated for 45 min at 37 °C. Wells were then extensively washed with PBS to remove non-adherent bacteria, dried at 60 °C for 10 mins, and stained with 0.1% crystal violet (Becton Dickinson) for 20 min at room temperature. After washing with water, 50% ethanol was added to solubilize the stain, and absorbance was measured at 600 nm.

### Red Blood Cell Agglutination Assay

To assess the effect of fluidic shear on *Escherichia coli* adhesion, we performed red blood cell (RBC) agglutination assays under both static and dynamic conditions using guinea pig RBCs (Colorado Serum Company, Inc., Denver, CO). For static conditions, a standard RBC rosette assay was conducted. Serial dilutions of *E. coli* suspensions (starting at OD_540_ = 2.0) were prepared in phosphate-buffered saline (PBS). A total of 50 μL of each bacterial dilution was mixed with 50 μL of a 1% RBC suspension in a U-bottom 96-well microtiter plate pre-coated with bovine serum albumin (BSA) to prevent nonspecific binding. Plates were incubated at room temperature without agitation for 60 minutes, after which images were taken. For dynamic conditions, a modified rocking assay was used. Equal volumes (50 μL each) of *E. coli* suspension (OD_540_ = 2.0) and 1% RBC suspension were combined on a BSA-coated concave serological glass plate. The plate was continuously rocked at room temperature to simulate shear forces, and images were taken after 20 minutes to assess RBC agglutination.

## Data and Code Availability

The data and Jupyter Notebooks for the CSD and BLI analyses are available at https://www.github.com/tuttlelm/FimH_L34K_paper.

## ACKNOWLEDGEMENTS

We thank Angelo Ramos for assistance with protein production and Natalie Stone for support with the collection and processing of HDX-MS data. We are also grateful to Anna Manchenko for assistance with RBC data collection and to Dagmara Kisiela for support with the mAb21 binding experiments. BLI data were collected at the Hans Neurath Biophysical Core at the University of Washington. HDX-MS data were acquired at the University of Washington’s Proteomics Resource (UWPR95794). Computational simulations were performed on the Expanse supercomputer at the San Diego Supercomputing Center through an ACCESS allocation (grant number TG-BIO240027), supported by the National Science Foundation. We are grateful to Peter Brzovic, Nicole Raniszewski, and Ronald Stenkamp for valuable discussions and for critically reading the manuscript. This work was supported by NIH grants K99GM141364 (to P.M.), R01AI171570 (to R.E.K. and E.V.S.), R01HL153253 (to G.I.), and R01AI119675 (to E.V.S. and W.E.T.).

## AUTHOR CONTRIBUTIONS

P.M., E.V.S., and R.E.K. designed the study. P.M. collected and analyzed data and wrote the manuscript. R.E.K. contributed to data analysis, wrote the manuscript and supervised the project. L.M.T. developed analytical tools and performed data analysis. G.I. conducted molecular dynamics simulations. L.A.C. and M.Y.M. carried out the AFM experiments. M.K.J performed data analysis for NMR and BLI. W.E.T. analyzed AFM and BLI data. E.V.S. collected and analyzed RBC data and carried out the mAb21 binding experiments.

