## Supplemental Figures and Tables for "Ultra-slow conformational dynamics and catch bond formation of a Bacterial Adhesin revealed by a single-domain variant of FimH"

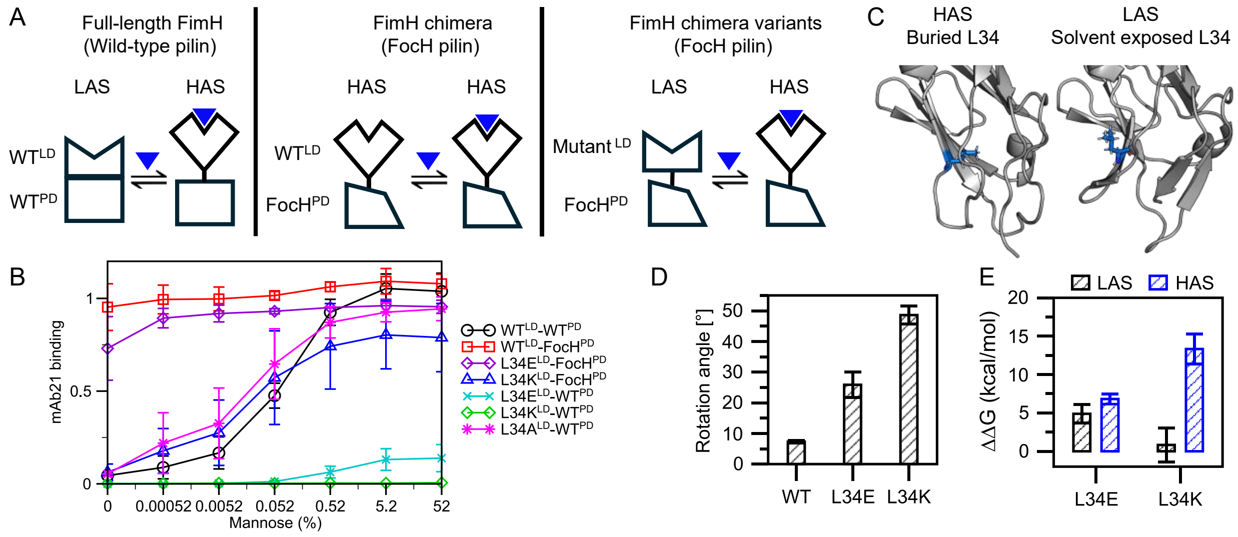

**Figure S1.** Toggle switch residue Leu34 substitutions in full-length FimH have differing effects on the ability to adopt the HAS conformation. (A) Schematic representations of the FimH variants and chimeras used in this study. Variants are referred to by their LD type (WT or a mutant) and their PD type (WT or FocH). The cartoons illustrate the expected conformational state of each, based on previous studies. (B) ELISA binding assays of HAS-specific monoclonal antibody, mAb21, conducted with variants of FimH as a function of increasing mannose concentrations. Constructs containing WT$^{LD}$, and L34A$^{LD}$ with WT$^{PD}$ exhibit mAb21 binding (and therefore, HAS conformation) at increasing mannose concentrations (black), but neither L34E$^{LD}$ (green) nor L34K$^{LD}$ (cyan) with the WT$^{PD}$ exhibit substantial binding, indicating they do not adopt HAS even when mannose is present. Chimeras of L34E$^{LD}$ or L34K$^{LD}$ with the incompatible FocH$^{PD}$ exhibit binding, but with different behaviors: 1) WT$^{LD}$-FocH$^{PD}$ and L34E$^{LD}$- FocH$^{PD}$ bind mAb21 in a mannose-independent manner, signifying these constructs predominantly adopt the HAS under all conditions; 2) L34K$^{LD}$-FocH$^{PD}$ exhibits a binding curve similar to the WT$^{LD}$-WT$^{PD}$ construct, signifying that the LD adopts the HAS in this context. (C) PyMOL models showing the position of the L34 side chain in HAS (PDB: 1UWF) where it is core-buried, and LAS (PDB: 4XOE) where it is solvent-exposed. (D) The sidechain rotation angle was determined by calculating the difference between the C$\alpha$-C$\beta$ vector of residue 34 during a simulation and the initial conformation. Reported are averages over two simulations while error bars denote standard errors of the mean (n=2). For each run, the first 10 ns of a 50-ns long simulation were discarded. (E) Free energy calculations indicate that the L34K substitution induces greater destabilization of the HAS conformation compared to L34E. Error bars show the standard errors of the mean of two simulations (n=3).

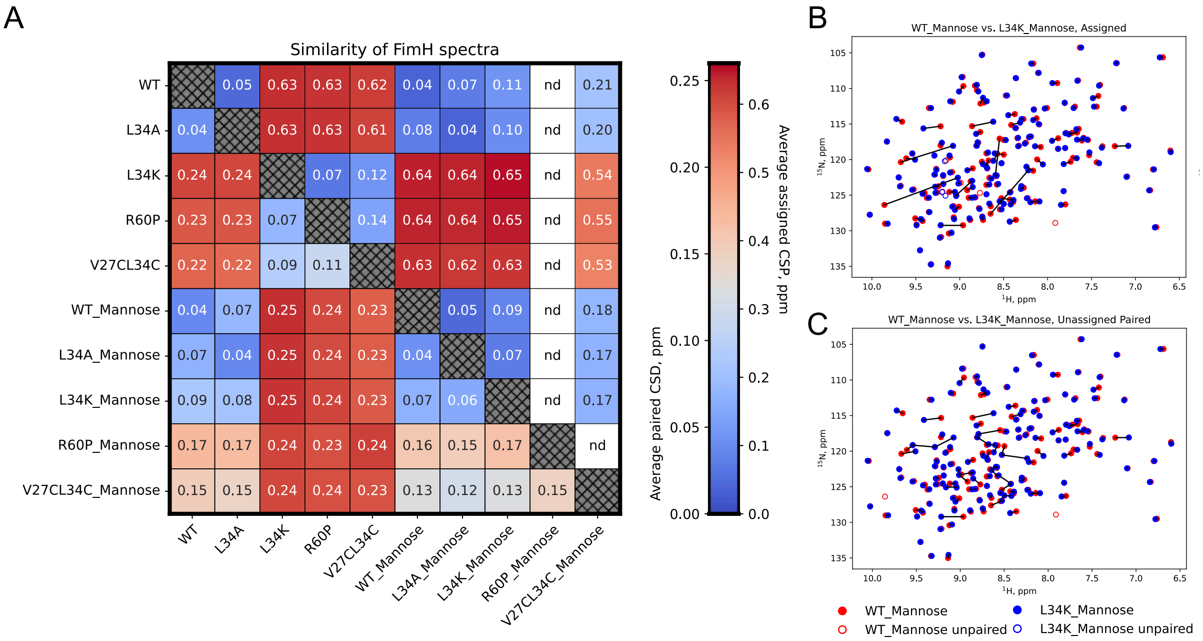

| | WT | L34A | L34K | R60P | V27CL34C | WT_Mannose | L34A_Mannose | L34K_Mannose | R60P_Mannose | V27CL34C_Mannose |
|---|---|---|---|---|---|---|---|---|---|---|
| WT | | 0.05 | 0.63 | 0.63 | 0.62 | 0.04 | 0.07 | 0.11 | nd | 0.21 |
| L34A | 0.04 | | 0.63 | 0.63 | 0.61 | 0.08 | 0.04 | 0.10 | nd | 0.20 |
| L34K | 0.24 | 0.24 | | 0.07 | 0.12 | 0.64 | 0.64 | 0.65 | nd | 0.54 |
| R60P | 0.23 | 0.23 | 0.07 | | 0.14 | 0.64 | 0.64 | 0.65 | nd | 0.55 |
| V27CL34C | 0.22 | 0.22 | 0.09 | 0.11 | | 0.63 | 0.62 | 0.63 | nd | 0.53 |
| WT_Mannose | 0.04 | 0.07 | 0.25 | 0.24 | 0.23 | | 0.05 | 0.09 | nd | 0.18 |
| L34A_Mannose | 0.07 | 0.04 | 0.25 | 0.24 | 0.23 | 0.04 | | 0.07 | nd | 0.17 |
| L34K_Mannose | 0.09 | 0.08 | 0.25 | 0.24 | 0.23 | 0.07 | 0.06 | | nd | 0.17 |
| R60P_Mannose | 0.17 | 0.17 | 0.24 | 0.23 | 0.24 | 0.16 | 0.15 | 0.17 | | nd |
| V27CL34C_Mannose | 0.15 | 0.15 | 0.24 | 0.24 | 0.23 | 0.13 | 0.12 | 0.13 | 0.15 | |

**Figure S2.** Assessment of spectral similarity for FimH variants. (A) Pairwise comparison of spectral similarity using the CSD score (below diagonal) or assigned CSP (above diagonal; nd, not determined) reveals the level of similarity between the LD variants in the presence and absence of mannose. (B,C) Comparison of paired peaks using the (B) assigned chemical shift or (C) unassigned paired chemical shift distance (CSD) analysis for mannose bound $WT^{LD}$ and $L34K^{LD}$. These are simulated spectra based only on the $^{1}H$ and $^{15}N$ peak positions. In the assigned analysis, open circles indicate a residue that is missing an assignment in one or both of the compared species. In the CSD analysis, open circles indicate peaks that were unpaired due to a difference in number of peaks in each spectrum. The corresponding CSP and CSD scores are 0.09 and 0.07 ppm, respectively, indicating a high level of similarity.

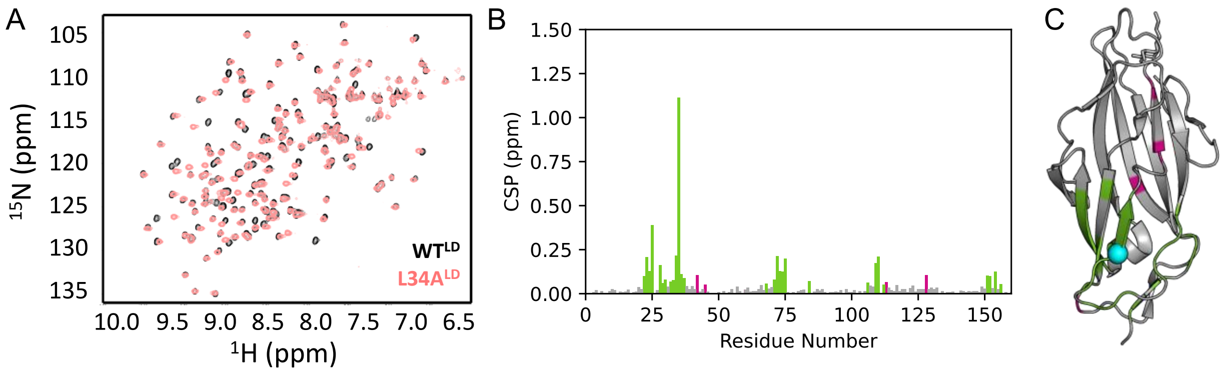

**Figure S3.** L34A$^{LD}$ adopts the HAS. (A) Overlay of the ($^{1}$H,$^{15}$N)-HSQC spectra of WT$^{LD}$ (black) and L34A$^{LD}$ (pink) in the absence of mannose shows extensive overlap between their chemical shifts. (B) The magnitude of CSPs between WT$^{LD}$ and L34A$^{LD}$, colored green or magenta for CSP > 0.05 ppm. (C) The significant chemical shift differences are localized to the site of mutation (residues within 10 Å of L34 are green), shown on PDB: 1UWF. Coloring is as shown in B, with L34 shown in cyan.

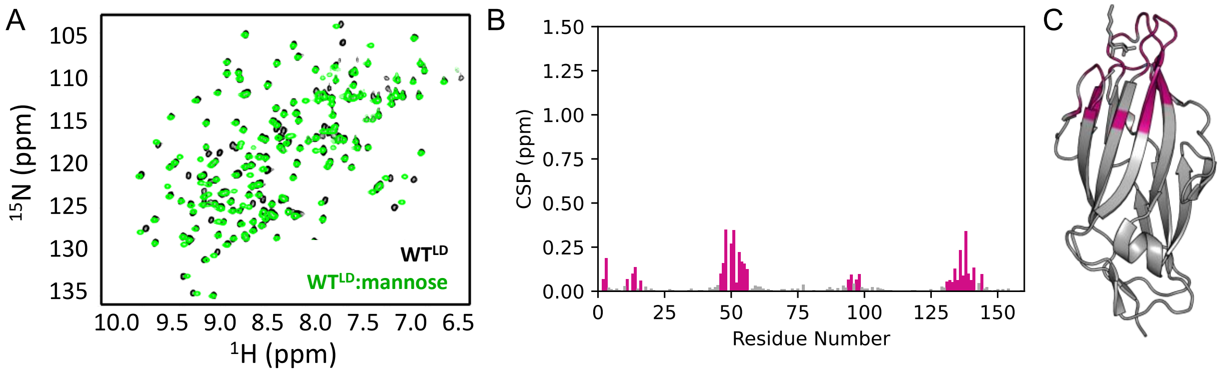

**Figure S4.** WT$^{LD}$ adopts HAS in the absence and presence of mannose. (A) ($^{1}$H,$^{15}$N)-HSQC spectra of WT$^{LD}$ (black) overlayed with WT$^{LD}$ in the presence of mannose (green) shows that ligand binding induces few chemical shift perturbations (CSPs) across the spectrum. (B) CSP plot shows that the magnitude of the CSPs in WT$^{LD}$ upon mannose binding is small (< 0.5 ppm), colored magenta for CSP > 0.05 ppm. (C) PyMOL plot (PDB:1UWF) shows that the CSPs > 0.05 ppm are localized in and around the mannose-binding pocket.

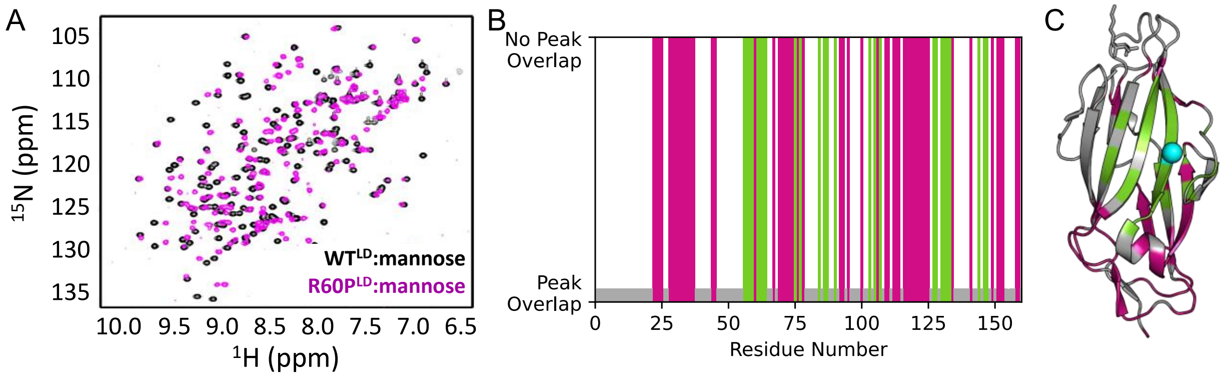

**Figure S5.** R60P$^{LD}$ does not convert to HAS and adopts a filled-intermediate LAS. (A) ($^{1}$H-$^{15}$N)-HSQC spectra of WT$^{LD}$ (black) and R60P$^{LD}$ (magenta) in the presence of mannose indicate that numerous chemical shift differences persist upon mannose binding. (B) The most pronounced differences, defined by the absence of peak overlap (no corresponding R60P$^{LD}$:mannose peak within 0.05 ppm of the assigned WT$^{LD}$:mannose peak; shown as tall bars) are clustered around the mutagenesis site (residues within 10 Å of R60 are green), with a secondary cluster of differences observed in the interdomain loops (magenta). (C) The "No Peak Overlap" residues are colored as in B on PDB: 1UWF; R60P is shown in cyan.

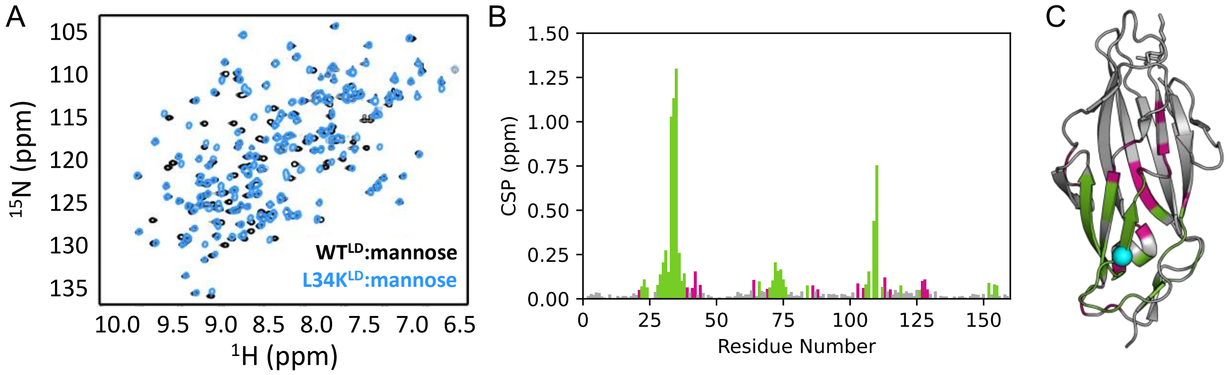

**Figure S6.** L34K$^{LD}$ adopts HAS in the presence of mannose. (A) ($^{1}$H-$^{15}$N)-HSQC overlays of WT$^{LD}$ and L34K$^{LD}$ in the presence of mannose. The spectra show significant overlap between the mannose-bound forms of WT$^{LD}$ and L34K$^{LD}$, indicating that L34K$^{LD}$ adopts the HAS upon mannose binding. (B) Magnitude of the CSP differences between mannose-bound WT$^{LD}$ and L34K$^{LD}$. (C) Mapping of the CSPs > 0.05 ppm on PDB: 1UWF reveals that they are localized near the L34K mutation site (residues within 10 Å of L34 are green). Coloring is as shown in B, with L34 shown in cyan.

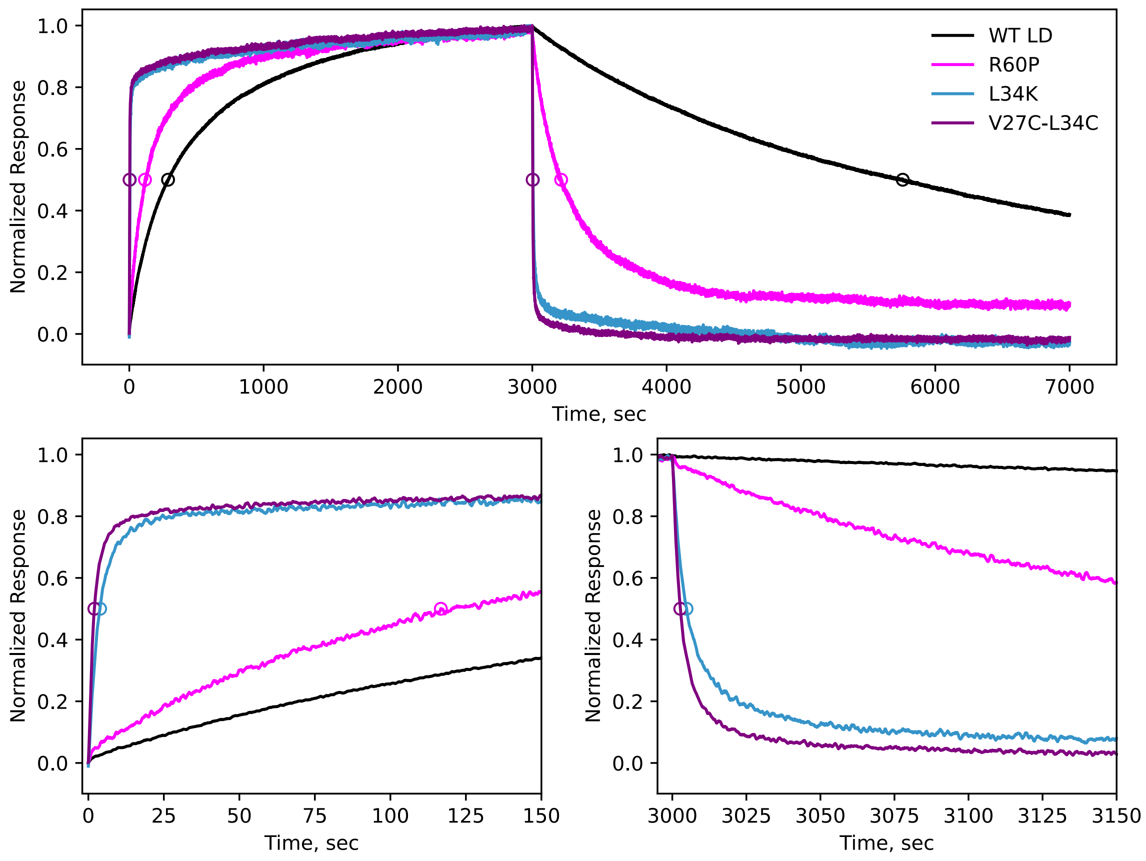

**Figure S7.** BLI response curves of LD variants interacting with immobilized mannose. Curves were collected at LD concentration of 5 μM. Circles are shown for the $t_{1/2,assoc}$ and $t_{1/2,dissoc}$ values for each variant, with bottom panels showing zoomed views. The LAS variants $L34K^{LD}$ and S-$S^{LD}$ (V27C-L34C) display a rapid initial association followed by a slower binding phase, and a rapid dissociation followed by a slower dissociation phase. $WT^{LD}$ displays slow association and dissociation. $R60P^{LD}$ shows behavior that is intermediate between the two extremes.

**A.**

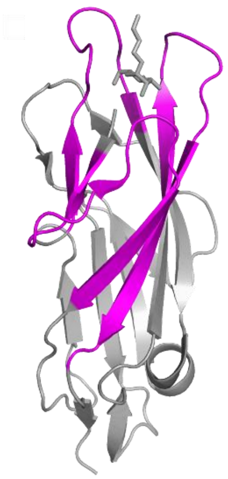

**Figure S8.** Deuterium uptake monitored by HDX-MS**.** A) The three regions showing differences in deuterium uptake after 30 minutes exchange—residues 3–21, 42–55, and 130–151—are highlighted in magenta. B) (below) Deuterium uptake plots for peptides in WT-empty, WT-filled, and L34K-filled HAS. Time points were collected at 5 s, 30 min, 1 hour, and 24 hours.

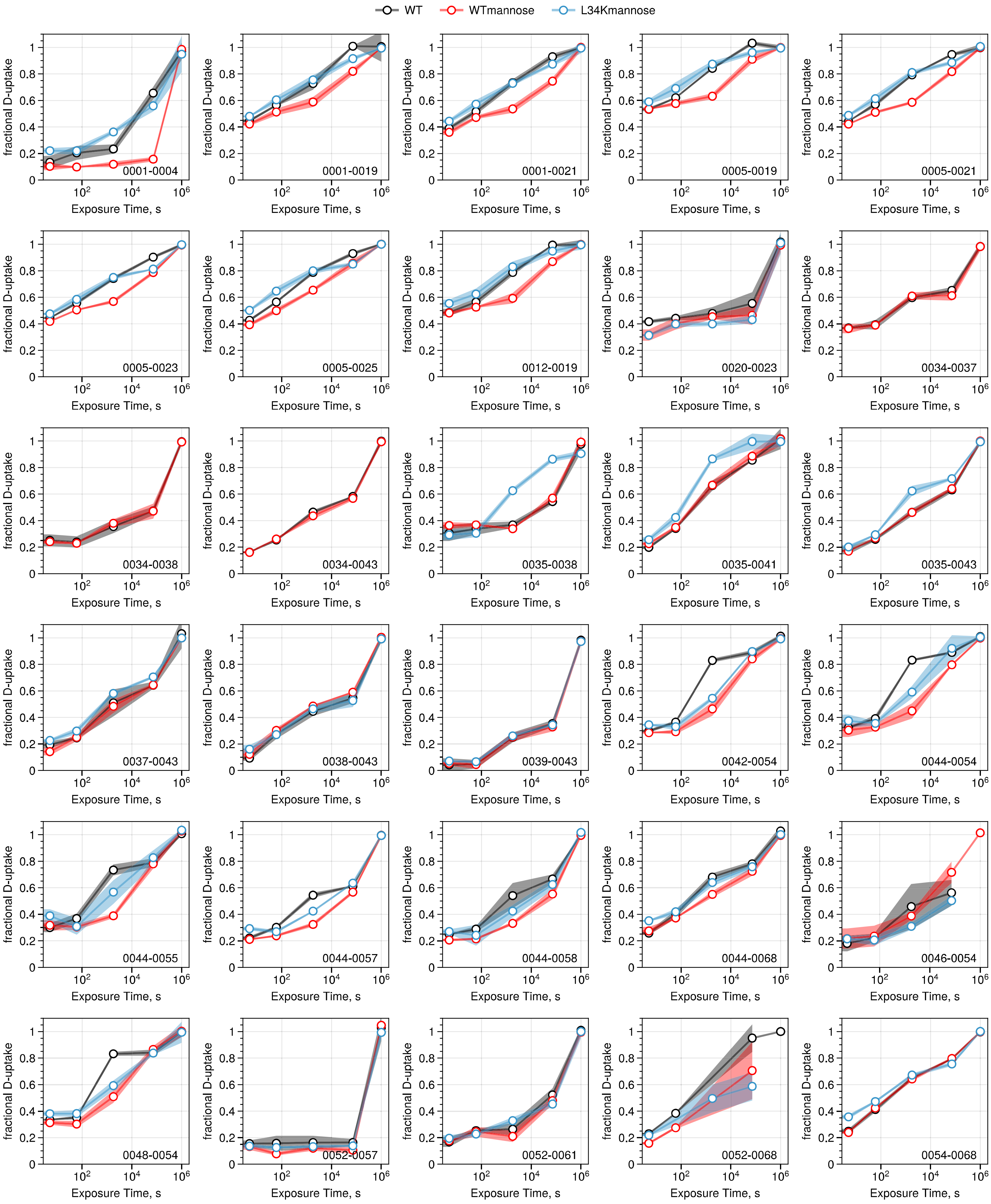

WT
WTmannose
L34Kmannose
fractional D-uptake
Exposure Time, s
0001-0004
0001-0019
0001-0021
0005-0019
0005-0021
0005-0023
0005-0025
0012-0019
0020-0023
0034-0037
0034-0038
0034-0043
0035-0038
0035-0041
0035-0043
0037-0043
0038-0043
0039-0043
0042-0054
0044-0054
0044-0055
0044-0057
0044-0058
0044-0068
0046-0054
0048-0054
0052-0057
0052-0061
0052-0068
0054-0068

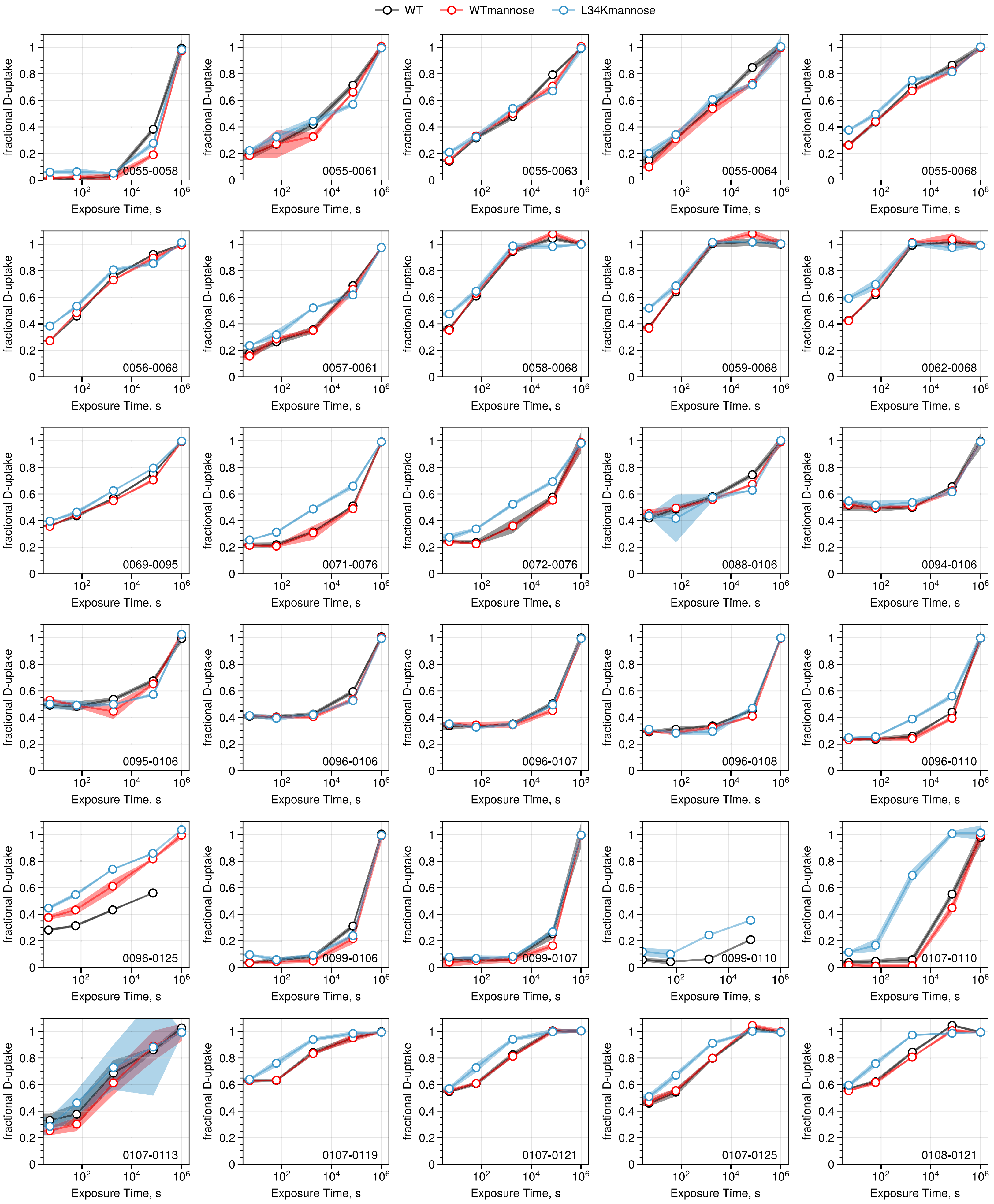

WT
WTmannose
L34Kmannose
fractional D-uptake
Exposure Time, s
0055-0058
0055-0061
0055-0063
0055-0064
0055-0068
0056-0068
0057-0061
0058-0068
0059-0068
0062-0068
0069-0095
0071-0076
0072-0076
0088-0106
0094-0106
0095-0106
0096-0106
0096-0107
0096-0108
0096-0110
0096-0125
0099-0106
0099-0107
0099-0110
0107-0110
0107-0113
0107-0119
0107-0121
0107-0125
0108-0121

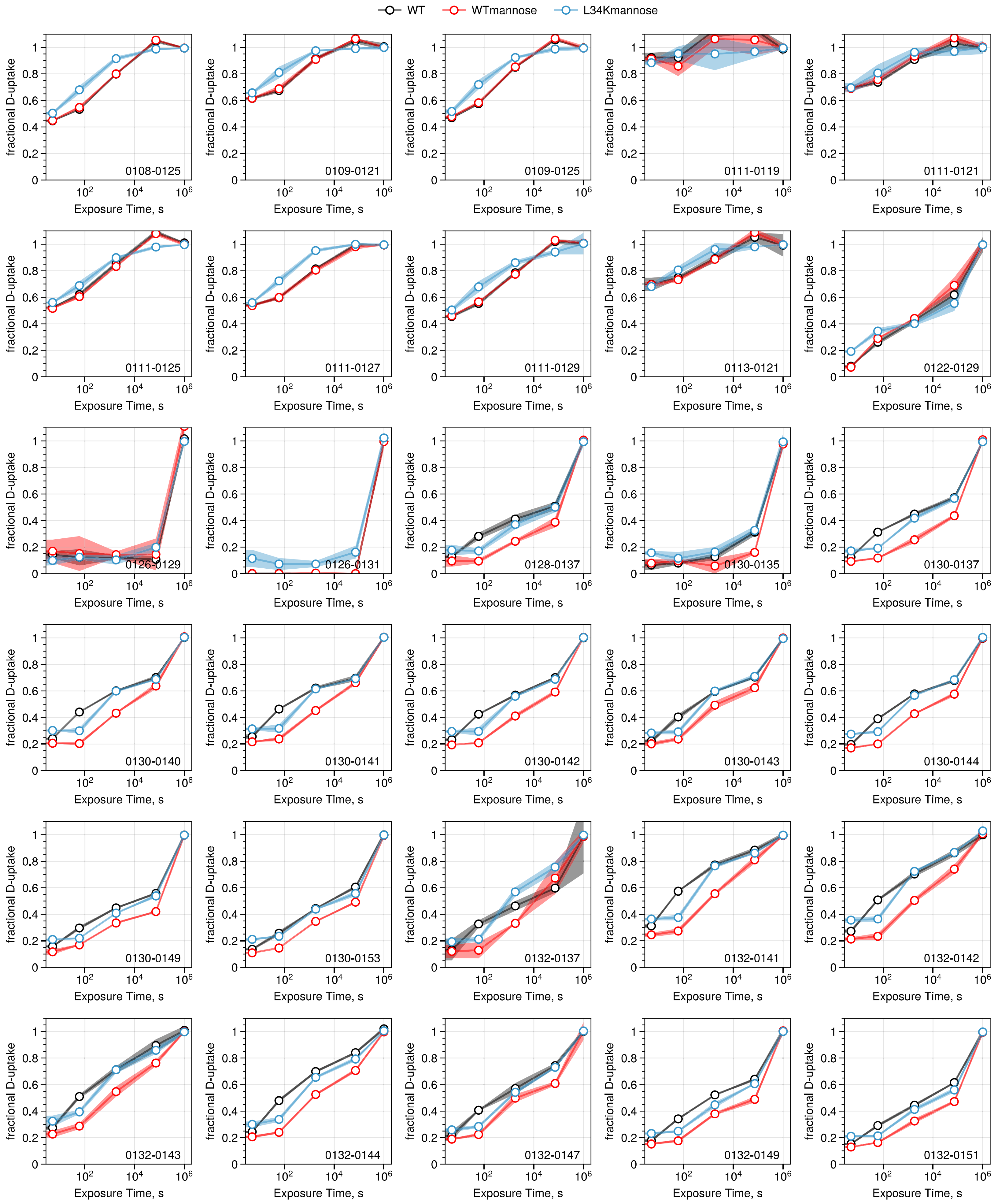
WT
WTmannose
L34Kmannose
fractional D-uptake
Exposure Time, s
0108-0125
0109-0121
0109-0125
0111-0119
0111-0121
0111-0125
0111-0127
0111-0129
0113-0121
0122-0129
0126-0129
0126-0131
0128-0137
0130-0135
0130-0137
0130-0140
0130-0141
0130-0142
0130-0143
0130-0144
0130-0149
0130-0153
0132-0137
0132-0141
0132-0142
0132-0143
0132-0144
0132-0147
0132-0149
0132-0151

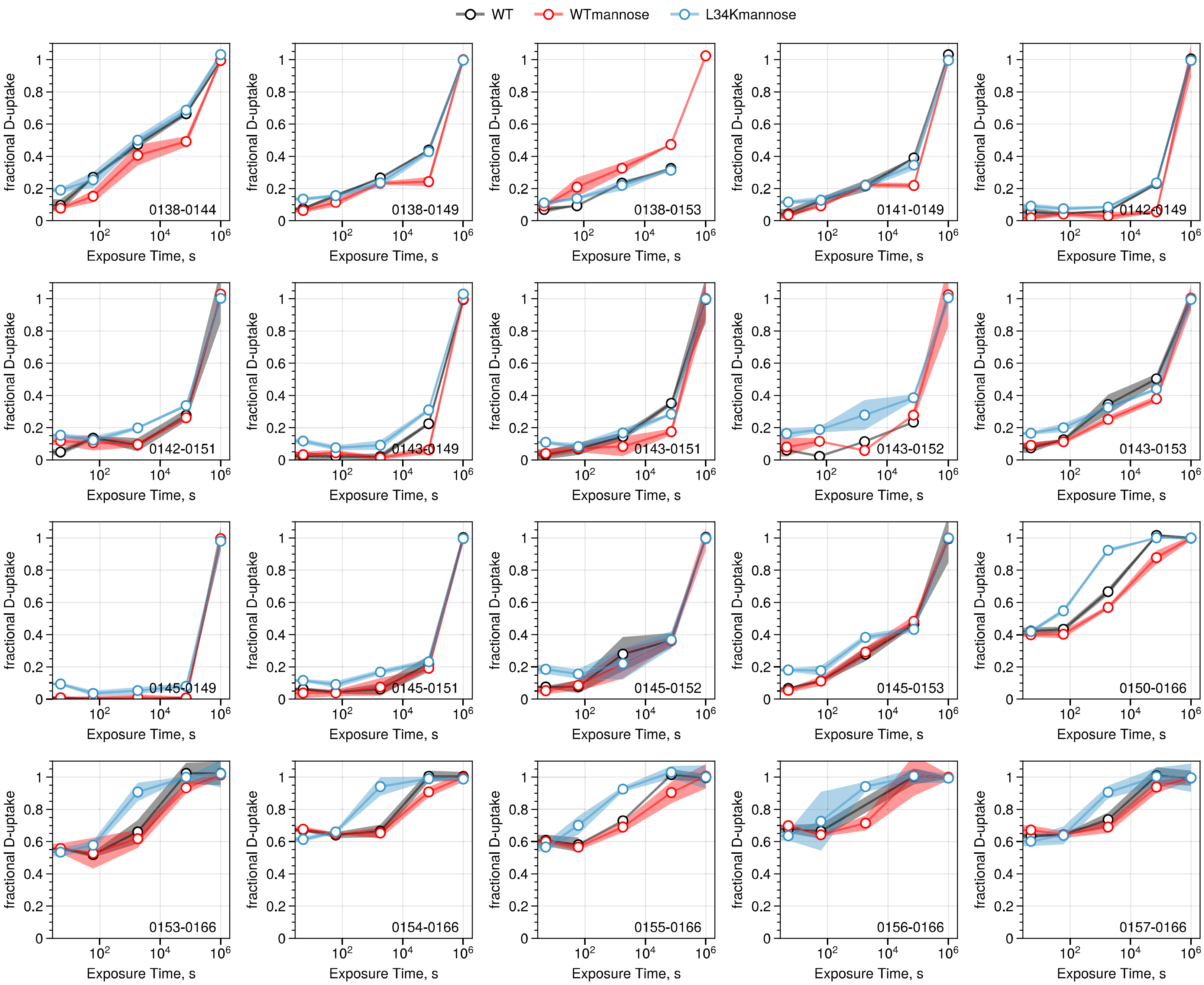

WT
WTmannose
L34Kmannose
fractional D-uptake
Exposure Time, s
0138-0144
0138-0149
0138-0153
0141-0149
0142-0149
0142-0151
0143-0149
0143-0151
0143-0152
0143-0153
0145-0149
0145-0151
0145-0152
0145-0153
0150-0166
0153-0166
0154-0166
0155-0166
0156-0166
0157-0166

**Figure S9.** Deuterium uptake monitored by HDX-NMR. Intensity decay curves and corresponding rate table for amide resonances are shown following exchange into $D_2O$. Faster decay indicates higher solvent accessibility; slower decay reflects more protected regions. Black traces are for the empty-HAS ($WT^{LD}$); red traces are for filled-HAS ($WT^{LD}$:mannose); blue traces are for L34K filled-HAS ($L34K^{LD}$:mannose). Y-axis is peak intensity; x-axis is time. Traces that are flat and have intensity near/at zero are fully exchanged at the first time point; traces that are flat with intensity > 0 have very slow deuterium exchange.

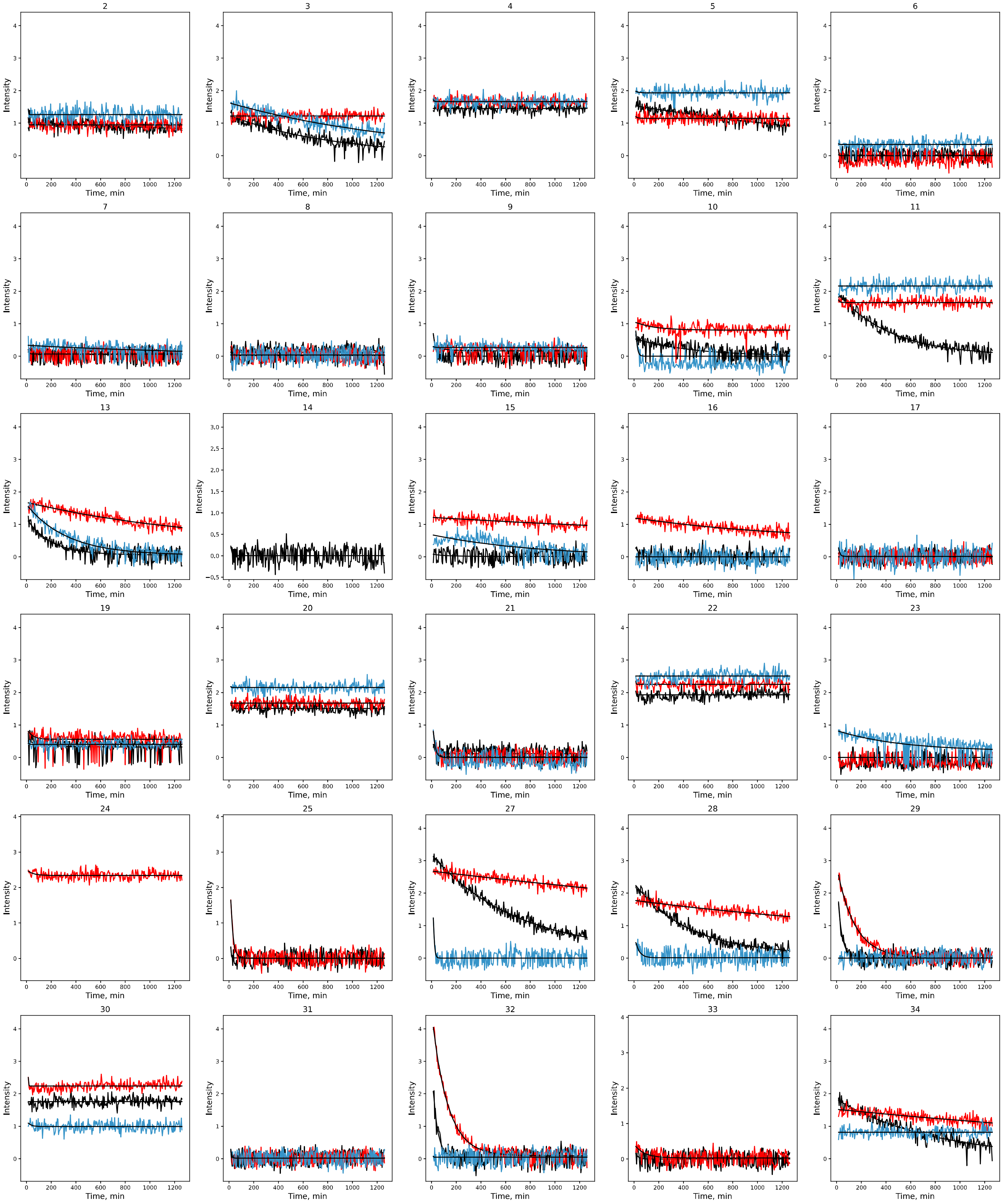
13
14
15
16
17
19
20
21
22
23
24
25
27
28
29
30
31
32
33
34
Intensity
Time, min

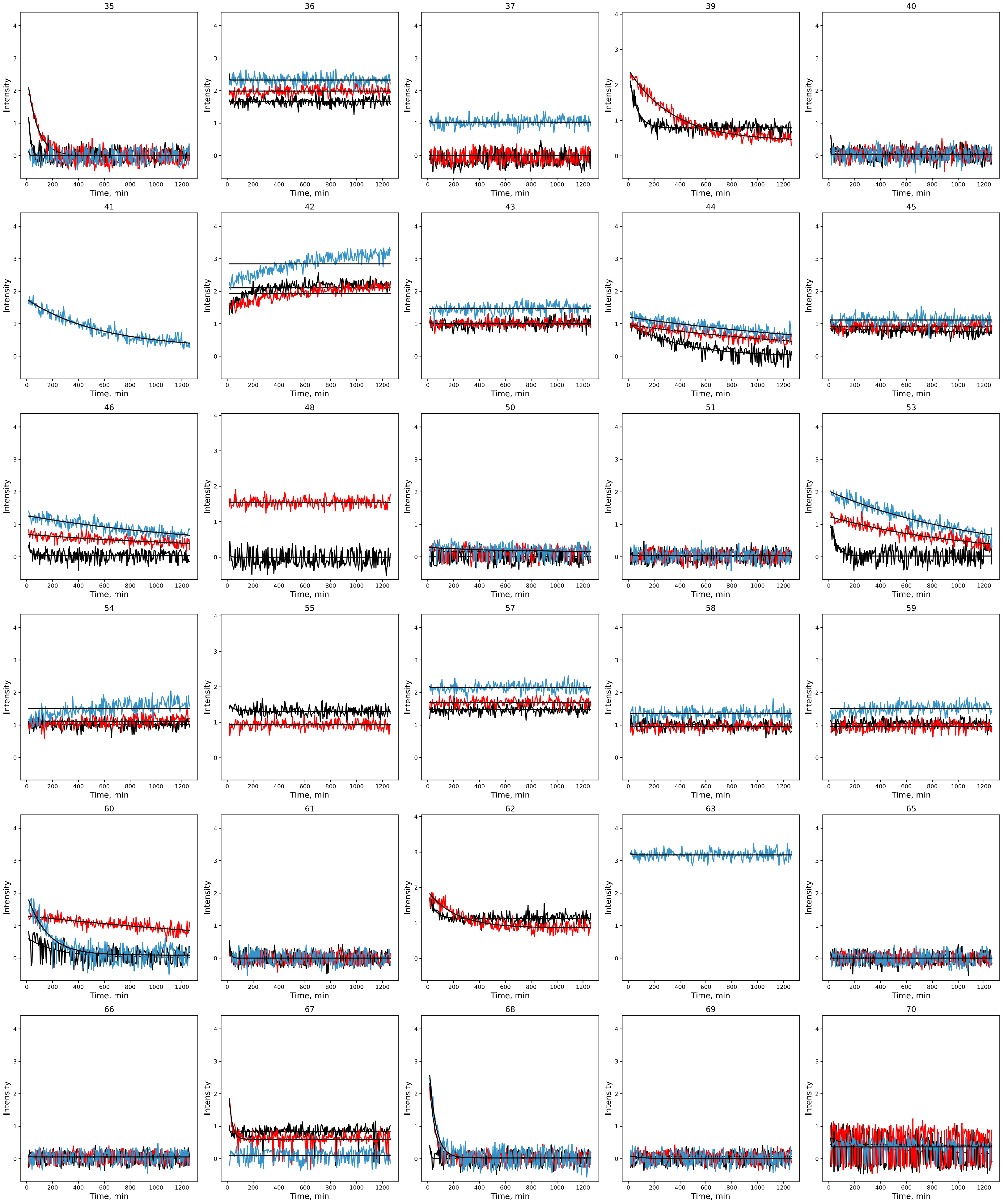
35
36
37
39
40
41
42
43
44
45
46
48
50
51
53
54
55
57
58
59
60
61
62
63
65
66
67
68
69
70
Intensity
Time, min

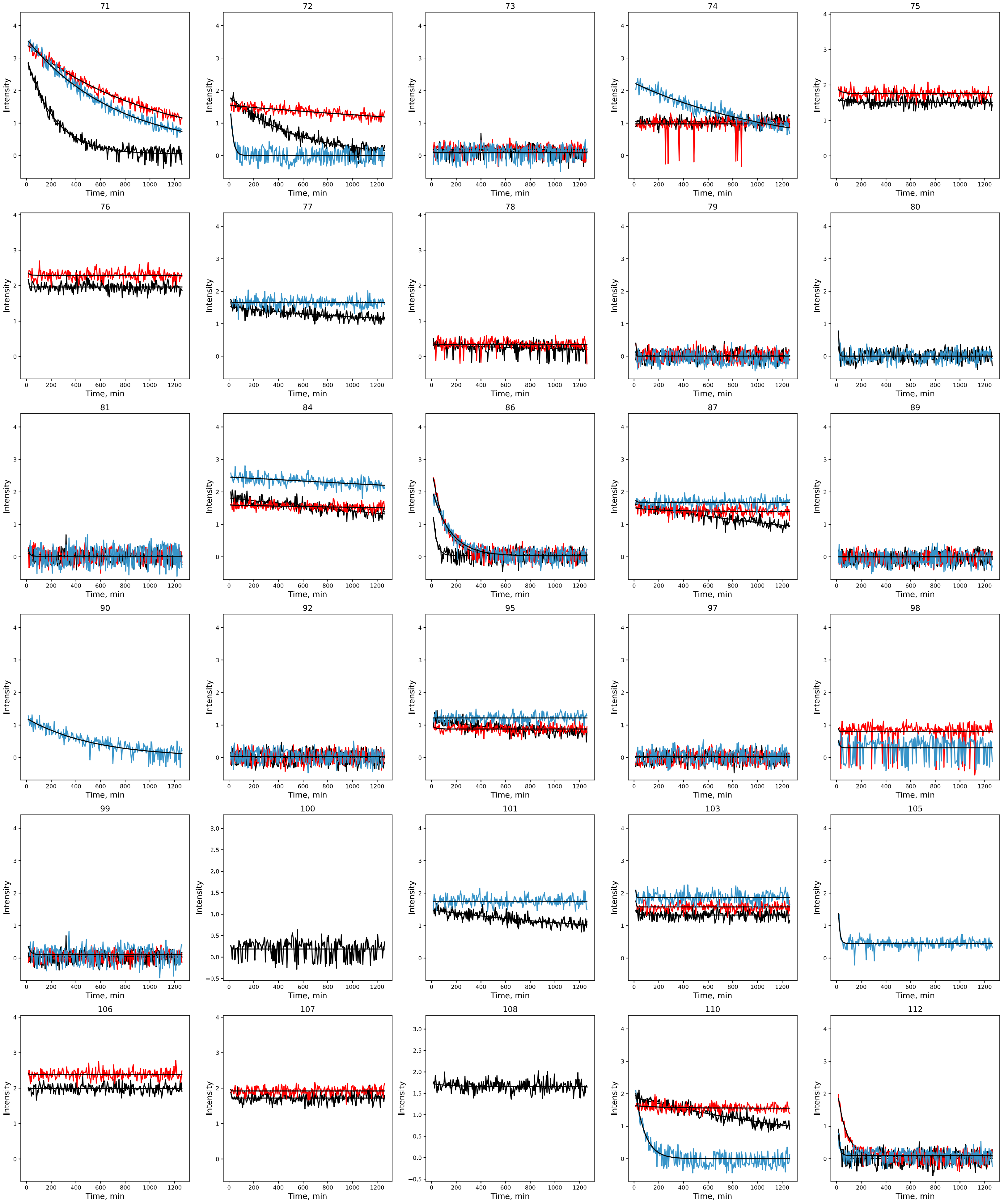
84
86
87
89
90
92
95
97
98
99
100
101
103
105
106
107
108
110
112
Intensity
Time, min

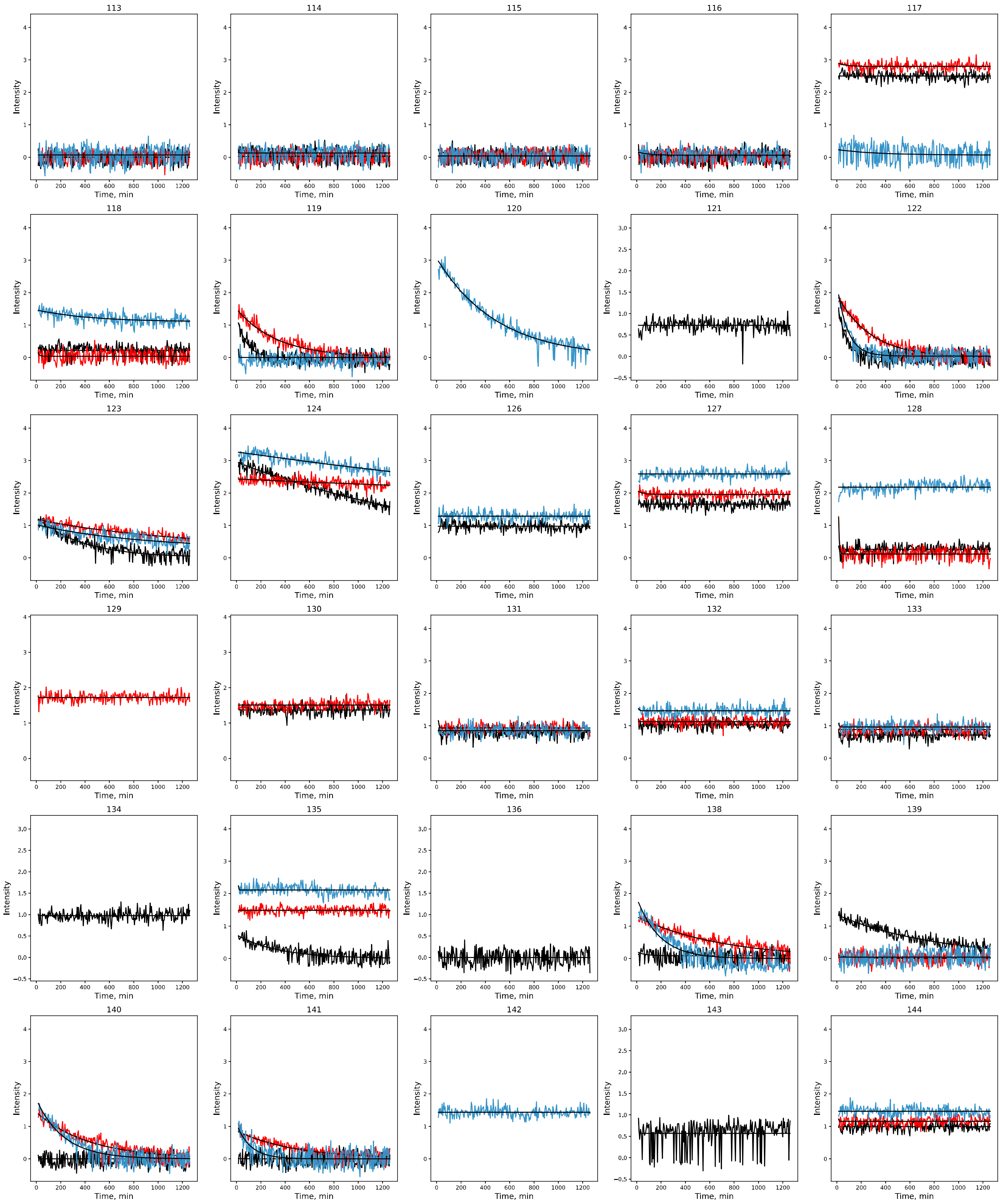
138
139
140
141
142
143
144
Intensity
Time, min

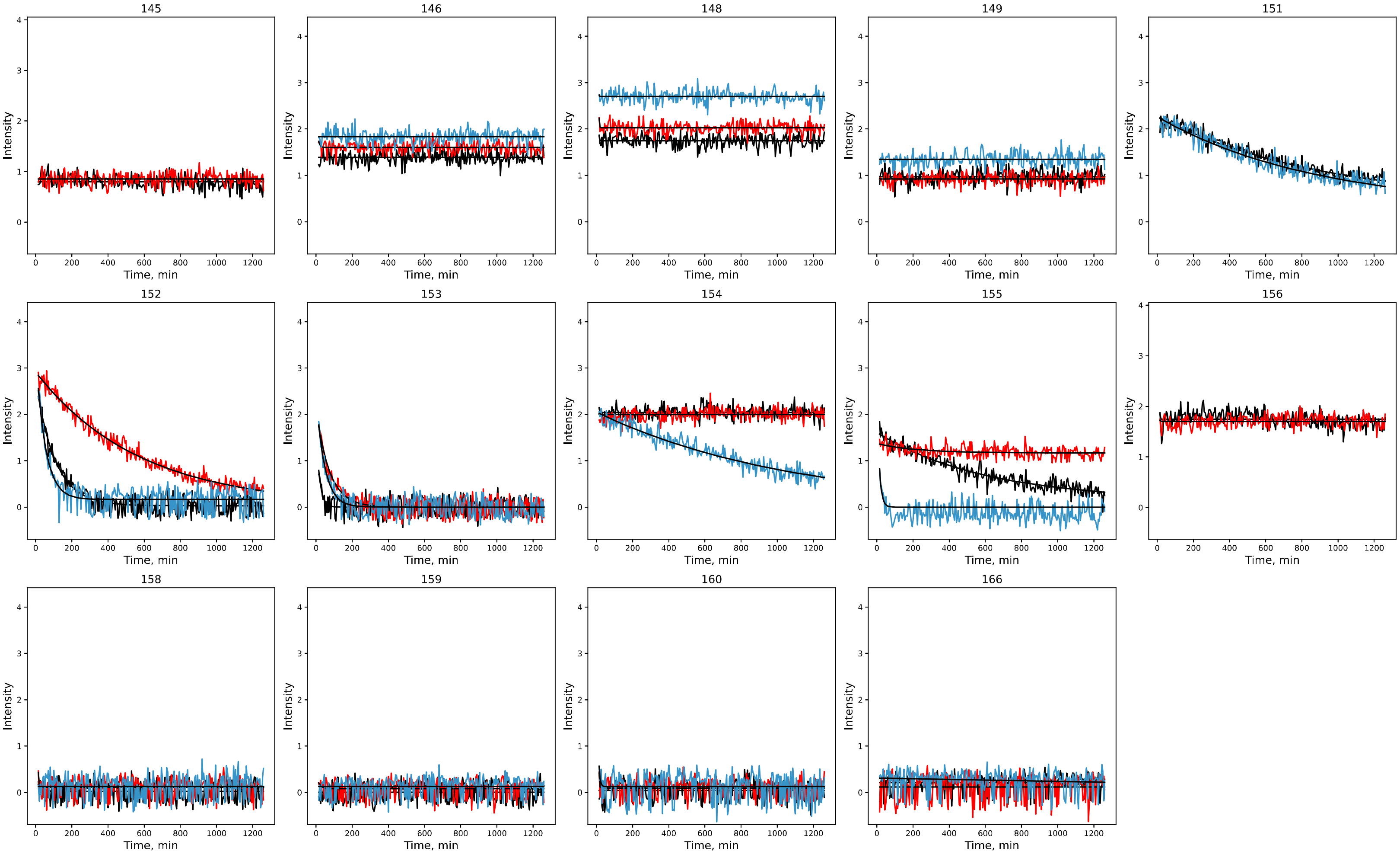
145
146
148
149
151
152
153
154
155
156
158
159
160
166
Intensity
Time, min

HDX-NMR rates ($min^{-1}$)

| Residue | WT | WT Mannose | L34K Mannose |
|---|---|---|---|
| 2 | slow | slow | slow |
| 3 | 1.21E-03 | slow | 6.73E-04 |
| 4 | slow | slow | slow |
| 5 | 4.15E-04 | slow | slow |
| 6 | fast | fast | fast |
| 7 | fast | fast | fast |
| 8 | fast | fast | fast |
| 9 | fast | fast | fast |
| 10 | fast | slow | fast |
| 11 | 2.32E-03 | slow | slow |
| 13 | 5.15E-03 | 6.57E-04 | 3.03E-03 |
| 14 | fast | overlap | overlap |
| 15 | fast | slow | fast |
| 16 | fast | 7.77E-04 | fast |
| 17 | fast | fast | fast |
| 18 | overlap | overlap | overlap |
| 19 | fast | slow | fast |
| 20 | slow | slow | slow |
| 21 | fast | fast | fast |
| 22 | slow | slow | slow |
| 23 | fast | fast | 1.94E-03 |
| 24 | overlap | slow | overlap |
| 25 | fast | 5.30E-02 | |
| 27 | 1.47E-03 | slow | fast |
| 28 | 2.26E-03 | slow | fast |
| 29 | 3.04E-02 | 6.84E-03 | fast |
| 30 | slow | slow | slow |
| 31 | fast | fast | fast |
| 32 | 2.57E-02 | 7.89E-03 | fast |
| 33 | fast | fast | overlap |
| 34 | 1.25E-03 | slow | slow |
| 35 | 6.06E-02 | 1.27E-02 | fast |
| 36 | slow | slow | slow |
| 37 | fast | fast | slow |
| 38 | overlap | overlap | overlap |
| 39 | 1.70E-02 | 2.91E-03 | overlap |
| 40 | fast | fast | fast |
| 41 | overlap | overlap | 1.76E-03 |
| 42 | slow | slow | slow |
| 43 | slow | slow | slow |
| 44 | 2.53E-03 | 5.79E-04 | 4.73E-04 |
| 45 | slow | slow | slow |
| 46 | fast | 4.93E-04 | 6.48E-04 |
| 47 | overlap | overlap | overlap |
| 48 | fast | slow | overlap |
| 50 | fast | fast | fast |
| 51 | fast | fast | fast |

| | | | |
|---|---|---|---|
| 52 | overlap | overlap | overlap |
| 53 | 2.19E-02 | 9.12E-04 | 8.68E-04 |
| 54 | slow | slow | slow |
| 55 | slow | slow | overlap |
| 56 | overlap | overlap | overlap |
| 57 | slow | slow | slow |
| 58 | slow | slow | slow |
| 59 | slow | slow | slow |
| 60 | 4.43E-03 | 3.34E-04 | 6.45E-03 |
| 61 | fast | fast | fast |
| 62 | 2.04E-02 | 4.56E-03 | overlap |
| 63 | overlap | overlap | slow |
| 64 | overlap | overlap | overlap |
| 65 | fast | fast | fast |
| 66 | fast | fast | fast |
| 67 | slow | 3.94E-02 | fast |
| 68 | fast | 2.36E-02 | 1.96E-02 |
| 69 | fast | fast | fast |
| 70 | fast | fast | fast |
| 71 | 4.44E-03 | 9.87E-04 | 1.36E-03 |
| 72 | 1.93E-03 | slow | 4.50E-02 |
| 73 | fast | fast | fast |
| 74 | slow | slow | 8.59E-04 |
| 75 | slow | slow | overlap |
| 76 | slow | slow | overlap |
| 77 | slow | overlap | slow |
| 78 | fast | fast | overlap |
| 79 | fast | fast | fast |
| 80 | fast | overlap | fast |
| 81 | fast | fast | fast |
| 84 | slow | slow | slow |
| 86 | 3.54E-02 | 9.48E-03 | 6.77E-03 |
| 87 | 4.92E-04 | slow | slow |
| 88 | overlap | overlap | overlap |
| 89 | fast | fast | fast |
| 90 | overlap | overlap | 1.83E-03 |
| 92 | fast | fast | fast |
| 93 | overlap | overlap | overlap |
| 94 | overlap | overlap | overlap |
| 95 | 1.52E-03 | slow | slow |
| 97 | fast | fast | fast |
| 98 | overlap | slow | fast |
| 99 | fast | fast | fast |
| 100 | fast | overlap | overlap |
| 101 | 6.56E-04 | overlap | slow |
| 103 | slow | slow | slow |
| 105 | overlap | overlap | 7.82E-02 |
| 106 | slow | slow | overlap |
| 107 | slow | slow | overlap |
| 108 | slow | overlap | |

| | | | |
|---|---|---|---|
| **109** | overlap | overlap | overlap |
| **110** | 5.67E-04 | slow | 1.29E-02 |
| **112** | fast | 1.31E-02 | fast |
| **113** | fast | fast | fast |
| **114** | fast | fast | fast |
| **115** | fast | fast | fast |
| **116** | fast | fast | fast |
| **117** | slow | slow | fast |
| **118** | fast | fast | slow |
| **119** | 1.25E-02 | 2.96E-03 | fast |
| **120** | overlap | overlap | 2.02E-03 |
| **121** | fast | overlap | overlap |
| **122** | 1.42E-02 | 3.61E-03 | 1.16E-02 |
| **123** | 2.62E-03 | 8.37E-04 | 1.19E-03 |
| **124** | 4.95E-04 | slow | slow |
| **125** | overlap | overlap | overlap |
| **126** | slow | overlap | slow |
| **127** | slow | slow | slow |
| **128** | fast | fast | slow |
| **129** | overlap | slow | overlap |
| **130** | slow | slow | overlap |
| **131** | slow | slow | slow |
| **132** | slow | slow | slow |
| **133** | slow | slow | slow |
| **134** | slow | overlap | overlap |
| **135** | 2.91E-03 | slow | slow |
| **136** | fast | overlap | overlap |
| **138** | fast | 1.43E-03 | 5.41E-03 |
| **139** | 1.19E-03 | fast | fast |
| **140** | fast | 2.29E-03 | 4.43E-03 |
| **141** | fast | 2.12E-03 | 8.78E-03 |
| **142** | overlap | overlap | slow |
| **143** | fast | overlap | overlap |
| **144** | slow | slow | slow |
| **145** | slow | slow | overlap |
| **146** | slow | slow | slow |
| **147** | overlap | overlap | overlap |
| **148** | slow | slow | slow |
| **149** | slow | slow | slow |
| **150** | overlap | overlap | overlap |
| **151** | 1.08E-03 | overlap | 1.14E-03 |
| **152** | 9.40E-03 | 1.74E-03 | 1.92E-02 |
| **153** | 5.36E-02 | 1.37E-02 | 2.11E-02 |
| **154** | slow | slow | 9.26E-04 |
| **155** | 1.63E-03 | slow | fast |
| **156** | slow | slow | overlap |
| **158** | fast | fast | fast |
| **159** | fast | fast | fast |
| **160** | fast | fast | fast |
| **166** | fast | fast | fast |

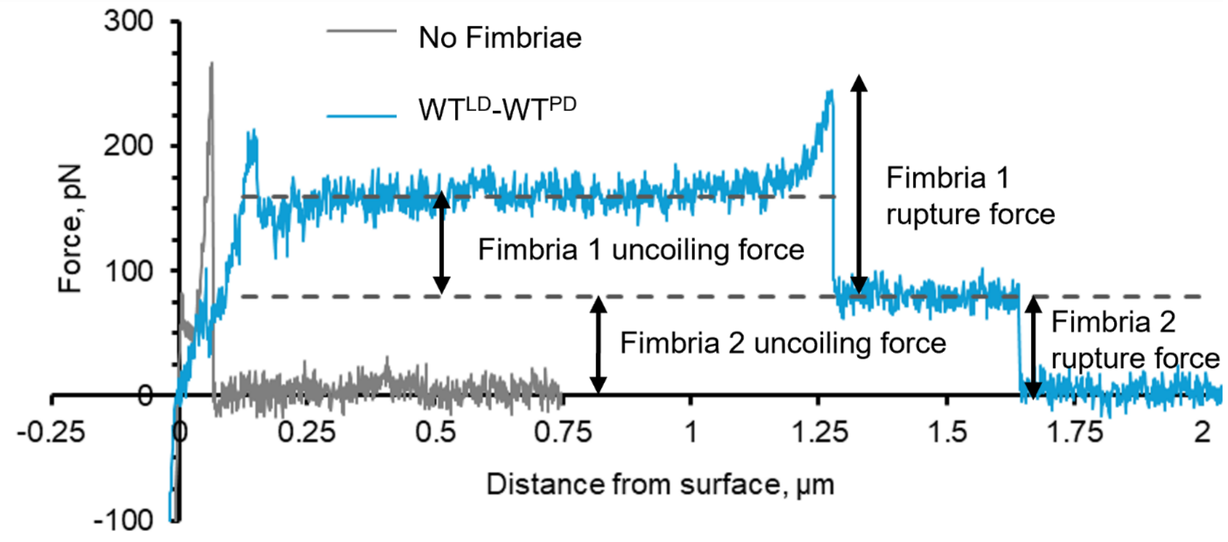

**Figure S10.** Sample data from Atomic Force Microscopy on fimbriae to illustrate how single FimH-mannose bond rupture forces are measured and distinguished from nonspecific adhesive interactions. Ruptures are defined as sharp significant drops in force (See methods for details.) The gray curve shows a control pull for a man-BSA coated cantilever pulling away after touching to a surface with no fimbriae, showing rupture of a nonspecific interaction that occurred within 0.25 µm of the surface. The blue curve shows sample force-extension curve for a man-BSA coated cantilever pulling away after touching to a surface coated with $WT^{LD}$-$WT^{PD}$ fimbriae. This curve shows three rupture events. The first was within 0.25 µm of the surface, so is within the distance threshold for ruptures that cannot be distinguished from nonspecific binding and therefore was not included in the analysis. The second, labeled "fimbria 1 rupture force" occurred while the force was ramping up as the first fimbria stretched after fully uncoiling at constant force. The third, labeled "fimbria 2 rupture force" occurred while the second fimbria was still uncoiling at constant force. Because fimbriae uncoil at around 75 pN, we would expect to see a significant number of ruptures to occur at this force because fimbriae experience the uncoiling force for more time than any other force and indeed this is seen in Figure 6C.

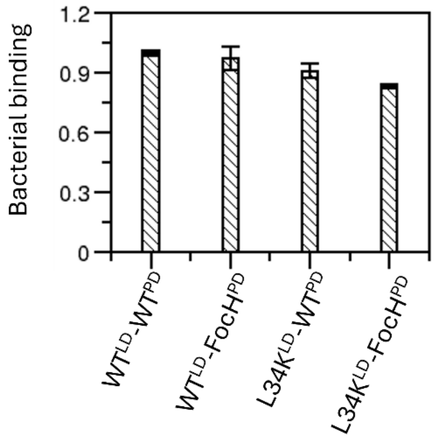

**Figure S11.** Bacterial binding to RNase B. Assessment of bacterial binding to RNase B, which contains high-mannose glycans (primarily $Man_5$), under static conditions. Binding was detected for each strain, confirming that functional FimH species are presented by each of the strains generated. Data are mean absorbance values at 600 nm ± SD.

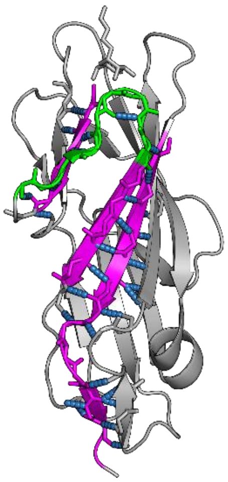

**Figure S12.** Interaction network flanking the Clamp loop. Residues flanking the clamp loop (green) in the LD include positions 1–5, which directly contact the mannose bound in the pocket, and residues 15–21, which form hydrogen bonds with the β-sheet spanning residues 143–157 that connects to the pilin domain (magenta). Hydrogen bonds are shown in blue. Structure shown from PDB entry 1UWF.

**Supplemental Table S1.** Half-life values from BLI association and dissociation phases for LD Variants. Values were calculated from the 5 μM LD response curves (left and middle columns) and the average from all curves was calculated for $t_{1/2,dissoc}$ (right column).

| **LD Variant** | **$t_{1/2,assoc}$ (sec) at [LD] = 5 μM** | **$t_{1/2,dissoc}$(sec) at [LD] = 5 μM** | **$\langle t_{1/2,dissoc} \rangle$ (sec) (S.D.)** |
|---|---|---|---|
| **$WT^{LD}$** | 288 | 2760 | 2717 (45) |
| **$L34K^{LD}$** | 3.90 | 4.80 | 5.2 (0.6) |
| **Ratio $WT^{LD}/L34K^{LD}$** | 74 | 575 | |

**Supplemental Table S2.** Summary of HDX/NMR behaviors.

| Species | # Fast | # Decay | # Slow | # Overlap |
|---|---|---|---|---|
| WT(empty) | 48 | 37 | 38 | 21 |
| WT(filled) | 38 | 29 | 48 | 30 |
| L34K(filled) | 43 | 30 | 47 | 34 |

**Supplemental Table S3.** Comparison of exchange rate ratios between WT(filled), WT(empty), and L34K(filled) States

| | Ratio >$10^2$ | 10 <Ratio < $10^2$ | 5 < Ratio < 10 |
|---|---|---|---|
| WT(filled)/WT(empty) | 15, 16, 46, 48, 138 | 3*, 11,27*, 95, 135, 140, 141 | 5, 68, 87, 110*, 124 |
| WT(filled)/L34K(filled) | 15#, 16#, 27*#, 98*#, 110*, 117*#, 155*# | 10#, 28#, 72, 74*, 119#, 154* | 3*, 29#, 32# |
| * Slow exchanging in WT(filled) so ratio is a lowest estimate<br># Fast exchanging in L34K(filled), so ratio is a lowest estimate | | | |
